# A *Mycobacterium tuberculosis* effector protein attacks host innate immunity by acting as an unusual ubiquitinating enzyme

**DOI:** 10.1101/2020.11.05.370551

**Authors:** Jing Wang, Pupu Ge, Zehui Lei, Zhe Lu, Lihua Qiang, Qiyao Chai, Yong Zhang, Dongdong Zhao, Bingxi Li, Jiaqi Su, Ruchao Peng, Yu Pang, Yi Shi, George Fu Gao, Xiao-Bo Qiu, Cui Hua Liu

**Author notes:** J.W., P.G., and Z.L. contributed equally to this work. To whom correspondence may be addressed. (G.F.G.), (X.-B.Q.), or (C.H.L.).

## Abstract

Protein kinase G (PknG), a eukaryotic type serine-threonine protein kinase (STPK) in *Mycobacterium tuberculosis* (Mtb), is secreted into the cytosol of infected macrophages to promote intracellular survival of mycobacteria and has been considered as a promising therapeutic target for tuberculosis (TB) treatment. However, the molecular details of Mtb PknG-host intracellular interactions remain obscure. Here, we demonstrate that PknG serves as both the ubiquitin-activating enzyme (E1) and the ubiquitin ligase (E3) to promote ubiquitination and degradation of tumor necrosis factor receptor-associated factor 2 (TRAF2) and TGF–β-activated kinase 1 (TAK1), and thus inhibits the NF-κB-mediated host innate immune responses. Surprisingly, PknG promotes the attachment of ubiquitin (Ub) to ubiquitin-conjugating enzyme (E2) UbcH7 via an isopeptide bond (UbcH7 K82-Ub), instead of a usual C86-Ub thiol-ester bond, and then promotes the discharge of Ub from UbcH7 by acting as an isopeptidase before attaching Ub to its substrates TRAF2 and TAK1. These results demonstrate that Mtb PknG promotes ubiquitination of the key components of the host innate immunity by acting as an unusual ubiquitinating enzyme to suppress innate immunity. Our findings provide a potential TB treatment via targeting unconventional ubiquitinating activities of PknG.

**Significance:** *Mycobacterium tuberculosis* (Mtb) protein kinase G (PknG), which is critical for Mtb intracellular survival, is a promising target for tuberculosis (TB) treatment. However, the molecular mechanisms underlying PknG-host interactions remain largely unclear. Here we demonstrate that PknG serves as both the ubiquitin-activating enzyme and the ubiquitin ligase to promote the ubiquitination and degradation of tumor necrosis factor receptor-associated factor 2 (TRAF2) and TGF-β-activated kinase 1 (TAK1), thus inhibiting NF-κB signaling activation. PknG promotes the attachment of ubiquitin to ubiquitin-conjugating enzyme UbcH7 via an isopeptide bond, instead of a usual thiol-ester bond, and releases the ubiquitin from UbcH7 by acting as an isopeptidase. These findings provide important information for rational development of TB treatment via targeting unconventional ubiquitinating activity of PknG.

## Introduction

Tuberculosis (TB), which is caused by the pathogen *Mycobacterium tuberculosis* (Mtb), remains the leading cause of human mortality caused by a single infectious agent. It led to about 10 million new infections and more than 1.45 million deaths including 0.25 million deaths due to co-infection with human immunodeficiency virus (HIV) in 2018 (1). What makes the matter worse is the widespread of drug-resistant Mtb strains that renders the currently available TB drugs ineffective. Thus, there is an urgent need to develop new TB drugs based on novel strategies and targets that are effective for drug-resistant TB. As a typical intracellular pathogen, Mtb can persist in host macrophages due to its ability to manipulate host signaling pathways and cellular processes, such as innate immune signaling pathways and phagocytosis (2, 3). Therefore, the information about the molecular mechanisms underlying Mtb-host interactions could be invaluable in developing novel effective and selective TB therapies based on Mtb-host interfaces.

Mtb eukaryotic-type serine/threonine protein kinases (STPKs) (4), which are speculated to mediate cross-talk between mycobacteria and host macrophage signaling pathways, have become prime targets for the development of novel TB therapeutics. However, the exact host regulatory functions and molecular mechanisms of the STPKs remain largely unknown. Of the 11 Mtb STPKs, the mycobacterial serine/threonine protein kinase G (PknG) is of particular interest mainly because of its role in Mtb intracellular survival and pathogenicity, as well as the fact that it is a secreted effector protein that is easy to target (5-7). PknG promotes the intracellular survival of Mtb by inhibiting the process of host phagosomal maturation (8). In addition, it was shown to regulate glutamate metabolism and to promote intrinsic antibiotic resistance, stress response and biofilm formation in mycobacteria (9-11). Indeed, blocking PknG kinase activity by the tetrahydrobenzothiophene inhibitor AX20017 resulted in enhanced clearance of the intracellular mycobacteria (12). However, Mtb PknG shows relatively high amino acid sequence homology (37%) to human protein kinase C-α (PKC-α) (13, 14), suggesting potential off-target adverse effects of AX20017 inhibition. Therefore, it is important to elucidate the detailed molecular mechanisms underlying Mtb PknG-mediated host-pathogen interactions to identify distinct targeting regions in PknG to improve selectivity of TB treatment.

Protein ubiquitination, which involves attaching the highly conserved ubiquitin (Ub) to a protein target, is a versatile post-translational modification that regulates a variety of cellular functions in eukaryotes including host innate immune signaling (such as NF-κB signaling) during infection (15). The typical ubiquitination process occurs by covalent attachment of Ub to other proteins via an isopeptide bond typically catalyzed by three enzymes, including ubiquitin-activating enzyme (E1), ubiquitin-conjugating enzyme (E2), and ubiquitin ligase (E3). During this process, E1 couples ATP hydrolysis to form a thiol-ester bond between its active cysteine (Cys) site and the C-terminal glycine (Gly) of Ub. Ub is then transferred onto the Cys residue in the active site of E2, which cooperates with an E3 (16-18). Ubiquitination is a reversible process catalyzed by the deubiquitinases (DUB), which specifically cleaves the isopeptide bond between Ub and modified proteins. During infection, the ubiquitination process could be targeted by effector proteins secreted by intracellular pathogens to subvert host innate immunity, thus promoting pathogen intracellular survival and proliferation. For example, the SidE family effectors from *Legionella pneumophila* could function as E3s to ubiquitinate host proteins using NAD as the energy source in the absence of E1 and E2 enzymes (19), which ubiquitination modification could be reversed by another *L. pneumophila* effector protein SidJ that acts as a DUB (20). However, how Mtb effector proteins regulate host ubiquitination system remains largely unclear.

Surprisingly, in our efforts to explore the molecular mechanisms underlying Mtb PknG-host interactions, we discovered that Mtb PknG binds to the host E2 protein UbcH7 via a novel ubiquitin-like (Ubl) domain, and promotes the attachment of Ub to UbcH7 via an isopeptide bond (UbcH7 K82-Ub), instead of a usual C86-Ub thiol-ester bond, and then promotes the discharge of Ub from UbcH7 by acting as an isopeptidase before attaching Ub to host target proteins including tumor necrosis factor receptor associated factor 2 (TRAF2) and TGF-β-activated kinase 1 (TAK1) for ubiquitination and degradation by acting as an unconventional E3 enzyme, leading to inhibition of NF-κB signaling activation. Taken together, these results indicate that Mtb PknG functions as an unusual ubiquitinating enzyme to suppress host innate immunity. Our findings reveal novel insights into the intricate interactions between Mtb and its host, providing a potential TB treatment via targeting unconventional ubiquitinating activity of Mtb PknG.

## Results

### Mtb PknG interacts with E2 protein UbcH7 during mycobacterial infectio

With an aim to better understand the regulatory roles and to identify host targets of Mtb PknG during mycobacterial infection, we attempted to screen its interaction proteins in the host by the yeast two-hybrid assay, and found that the host E2 protein UbcH7 interacts directly with Mtb PknG (Fig. 1*A* and *SI Appendix*, Table S1). To verify this interaction, we deleted the gene encoding PknG (*pknG*) in *M. tuberculosis* strain H37Rv (Mtb *ΔpknG*) and challenged macrophage-like U937 cells with wild-type (WT) Mtb or Mtb *ΔpknG* strain for immunoprecipitation assay. The results showed that Mtb PknG interacted with UbcH7 in the infected macrophages (Fig.1*B*). To determine the direct and specific interactions of PknG with UbcH7, we also performed pull-down assays *in vitro*, and demonstrated that PknG had a strong interaction with UbcH7, but not with other E2s including UbcH5a or UbcH8 (Fig.1*C*). We then re-examined the structure of Mtb PknG (PDB ID: 2PZI) to identify their potential interacting motifs. There are at least 70 distinct ubiquitin-like (Ubl) families in eukaryotes, all of which contain a β-grasp fold (21). Notably, in addition to the conserved kinase domain (amino acids 151–396) and tetratricopeptide repeat (TPR) domain (amino acids 506–568), a Ubl domain as a potential E2 enzyme-binding domain was predicted at the N-terminal region of PknG (Fig. 1*D*). PknG contains a region (amino acids 140–234) similar to the β-grasp fold domain. Like that in Ub and the ubiquitin-fold domain (UFD) of E1 (Fig. 1*E*), this PknG region contains 5 β-strands and an α-helix (Fig. 1*F*). The orientation of 5 strands in PknG is highly similar to that of Ub or the UFD of E1, though its α-helix lies at a different orientation.

**Fig. 1.**
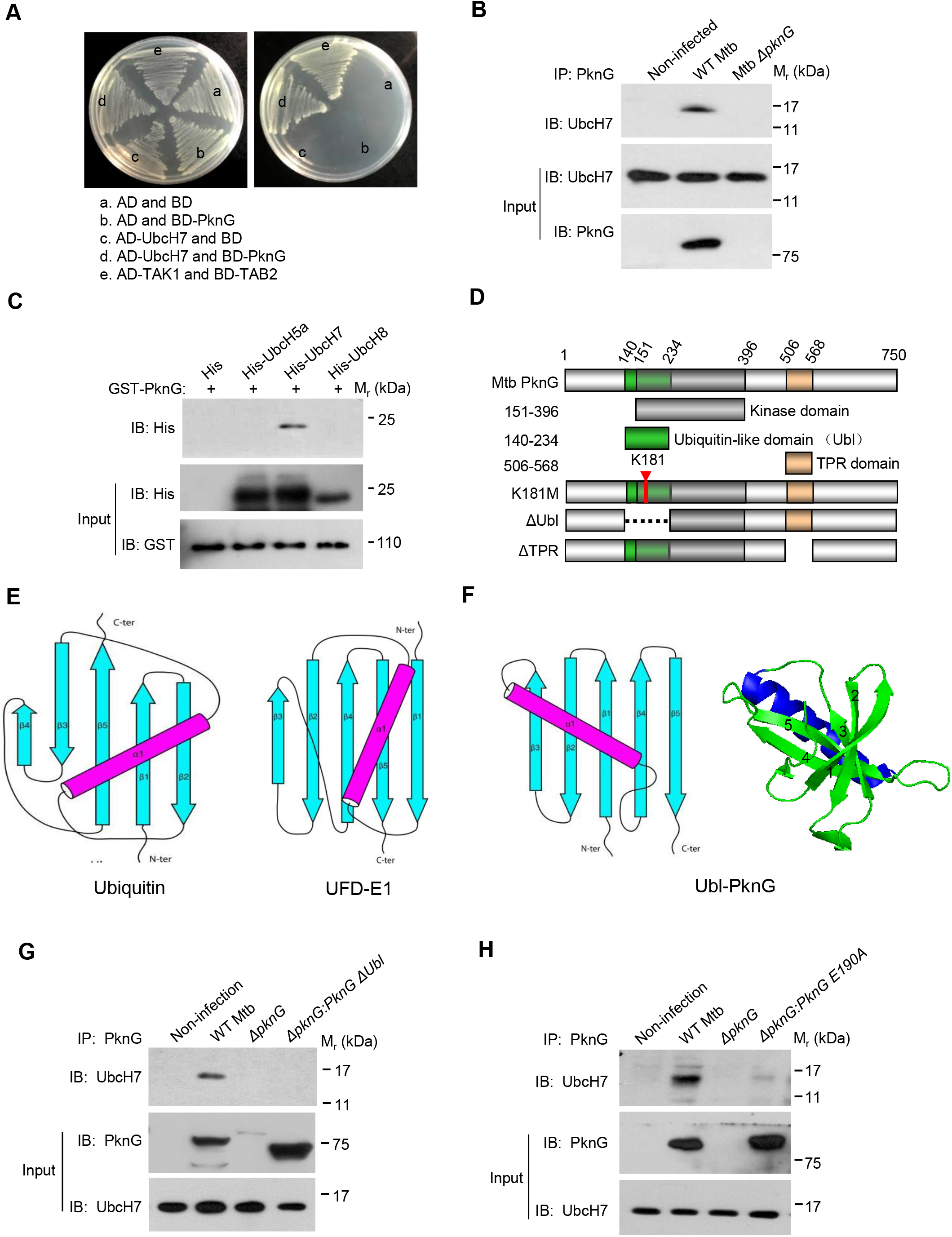
Mtb PknG interacts with E2 protein UbcH7 through Ubl domain. (*A*) Yeast two-hybrid assay for interaction of PknG with UbcH7. Yeast strains were transformed with the indicated plasmids in which TAK1-TAB2 interaction serves as a positive control. Left, low stringency. Right, high stringency. (*B*) Immunoprecipitation (IP) of UbcH7 by Mtb PknG in U937 cells. Cells were infected with wild-type (WT) Mtb or Mtb *ΔpknG* at an MOI of 1. Non-infected cells were used as control. After 4 h, cells were lysed and immunoprecipitated with antibody against PknG. (*C*) Pull-down of His-PknG (10 μg each) by GST-tagged E2s (5 μg each). (*D*) Schematic diagram of Mtb PknG domains. (*E-F*) Topology diagrams of the β-grasp folds of ubiquitin (*E, left*), the ubiquitin-fold domain (UFD) of the E1 enzyme (*E*, right), and the putative Ubl domain in PknG (*F*, left) are shown, and a cartoon depiction of β-grasp fold of PknG is included (*F*, right). Strands in topology diagrams are illustrated as cyan arrows, and helices are in magenta. (*G*) IP of UbcH7 by PknG or PknGΔUbl in U937 cells. Cells were infected with wild-type (WT), *ΔpknG, ΔpknG:pknG*, or *ΔpknG:pknGΔUbl* Mtb strain as in (*B*). *(H)* IP of UbcH7 by PknG or PknG E190A in U937 cells. Cells were infected with WT, *ΔpknG, ΔpknG:pknG*, or *ΔpknG:pknG E190A* Mtb strain as in (*B*).

Given the high diversity of β-grasp folds even in eukaryotes, we wondered whether the prokaryotic β-grasp fold-like region of PknG can serve as a ubiquitin-like domain. Because UbcH7 can bind the α-helix of Ub (13, 22, 23), we thus hypothesized that the α-helix (amino acids 188–204) in the β-grasp-fold-like region may be the potential binding region for the interaction of PknG with UbcH7 (*SI Appendix*, Fig. S1*A*). Indeed, this PknG region had a strong interaction with UbcH7, while deletion of its β-grasp-fold-like region abolished this interaction in the infected macrophages and transfected HEK293T cells (Fig. 1*G* and *SI Appendix*, Fig. S1*B*). We then did serial mutation screening in the potential UbcH7-interacing region of PknG through constructing a number of PknG mutants in which the residues covering the potential binding sites were mutated into alanine (Ala) (*SI Appendix*, Fig. S1*C*), and found that PknG Glu190Ala (PknG E190A) mutation did not affect the kinase activity of PknG (*SI Appendix*, Fig. S1*D*), but it abolished the interaction of PknG with UbcH7 (Fig.1 H and *SI Appendix*, Fig. S1*E*). Taken together, Mtb PknG interacts with host E2 protein UbcH7 via a novel Ubl domain during mycobacterial infection.

### Mtb PknG acts as an unconventional E1 with isopeptidase activity

The typical ubiquitination process requires a cascade of three enzymes including E1, E2, and E3 that activates, conjugates, and transfers Ub to the substrate sequentially (16). Normally, the E2 enzyme UbcH7 is charged with Ub by an E1 and subsequently interacts with E3s, *e*.*g*., Parkin and HHAIR, to mediate substrate ubiquitination (24). We thus sought to determine whether Mtb PknG interacts with UbcH7 to mimic a eukaryotic E1 or E3 during mycobacterial infection. We started by conducting the ubiquitin-conjugation assay for UbcH7, and found that PknG was able to function as an E1 to promote the loading of Ub onto UbcH7 using ATP as the source of energy, which is indicated by a molecular weight shift in SDS–PAGE (Fig. 2*A*). To exclude the potential interference of the kinase activity of PknG on its ubiquitination function, we performed liquid chromatography mass spectrometry (LC-MS) using PknG Lys181Met (K181M, a kinase-dead mutant) (7), and found that PknG K181M could hydrolyze ATP to release ADP (revised *SI Appendix*, Fig. S2*A*), which process is different from that mediated by the conventional E1 that usually hydrolyzes ATP to release AMP (25-28). Consistently, through conducting western blotting analysis using ATP analogs (including ATP-γ-S, AMPCPP, and AMPPNP) to replace ATP, we found that PknG could use AMPCPP and AMPPNP, but not ATP-γ-S, to catalyze Ub-conjugation onto UbcH7, suggesting that ATP is hydrolyzed at the γ site during the reaction (*SI Appendix*, Fig. S2*B*). Interestingly, unlike the classic Ub conjugation of E2s that require the E1 to form a thiol-ester bond between their active Cys and the C-terminal Gly of Ub, PknG could directly catalyze Ub-conjugation onto UbcH7 in a kinase activity-independent and Ubl domain-dependent manner (Fig. 2*B* and *SI Appendix*, Fig. S2*C*), bypassing the formation of PknG-Ub linkage step (*SI Appendix*, Fig. S2*D* and *E*). We then further did serial mutation screening through constructing a number of PknG truncations or mutants, and found that Pro57Ala mutation in the N-terminal region of PknG (PknG P57A) abolished its E1 activity, suggesting that Pro57 is required for the E1 activity of PknG towards UbcH7 (Fig.2C and *SI Appendix*, Fig. S2 *F–I*). In a canonical ubiquitination reaction, the charged E1∼Ub recruits cognate E2s, and Ub is then transferred from the catalytic Cys of E1 to the E2 catalytic Cys residue (16). We then sought to identify the residue of PknG that mediates UbcH7 ubiquitin conjugation. Unexpectedly, when the critical Cys residues of PknG were replaced by Ala, Ub conjugation of UbcH7 could still be easily detected, indicating that PknG might catalyze ubiquitin conjugation of UbcH7 independent of the formation of Cys∼Ub linkage (*SI Appendix*, Fig. S3*A*). We then performed mass spectrometry (MS) analysis to identify the Ub-binding sites of the charged UbcH7∼Ub mediated by PknG, and detected the identity of the Ub peptides with a GlyGly or LeuArgGlyGly modification on the residues of UbcH7. Specifically, we found that Lys82 was the new Ub modification site catalyzed by Mtb PknG (Fig. 2*D* and Dataset S1). We then further replaced Lys82 (a new Ub modification site catalyzed by PknG) or Cys86 (a canonical catalytic residue of the E2 precharged with Ub by E1) (16) with Ala in UbcH7 (UbcH7 K82A and C86A) for ubiquitin conjugation assay of UbcH7, and found that Lys82, but not Cys86, of UbcH7 was the crucial site for PknG-catalyzed ubiquitin conjugation (*SI Appendix*, Fig. S3*B*). Taken together, these results indicate that PknG exhibits Ubl domain-dependent unconventional E1 activity, which could directly catalyze Ub conjugation onto the Lys82 of UbcH7 in the presence of ATP, without forming an intermediate PknG∼Ub thiol-ester linkage.

**Fig. 2.**
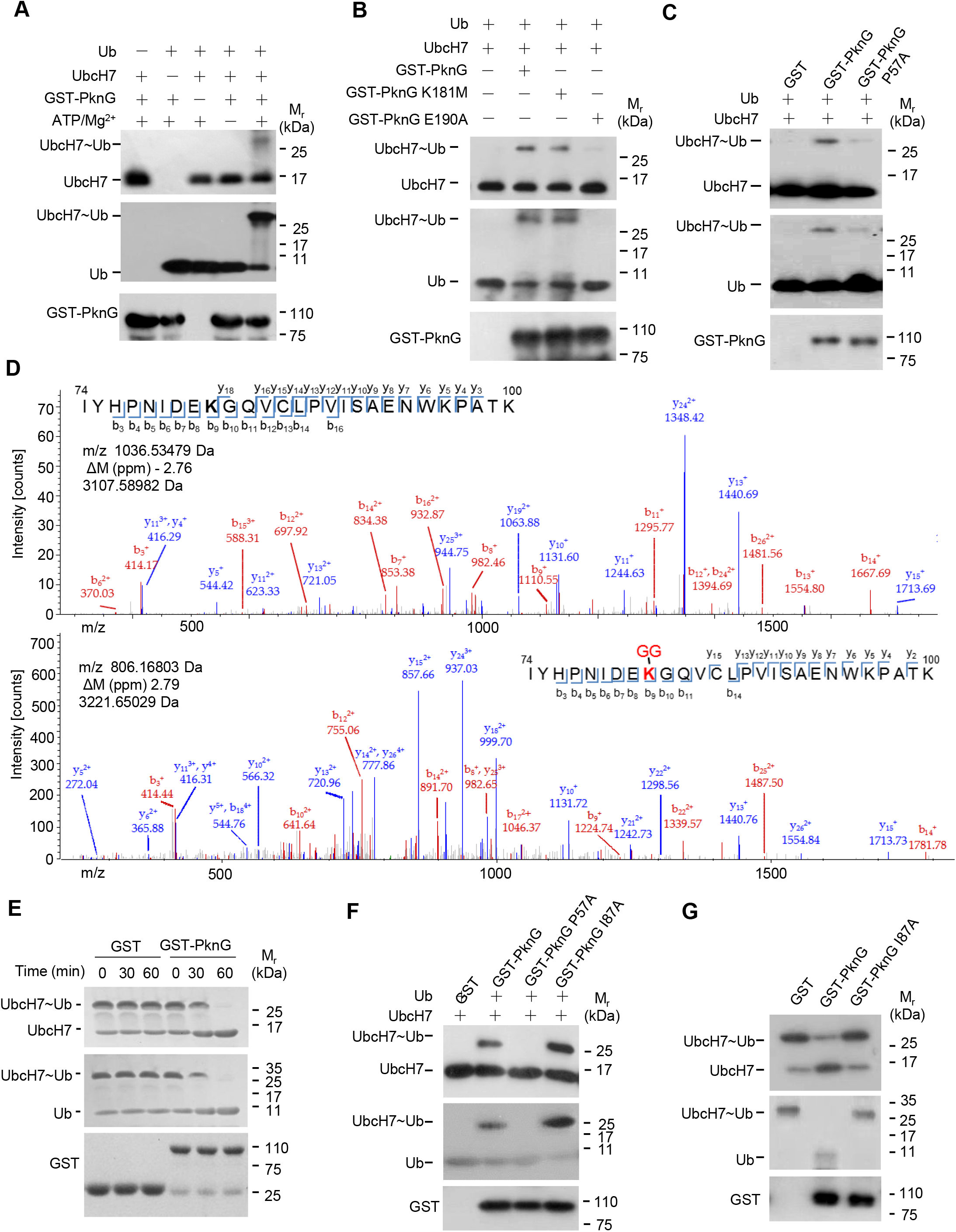
Mtb PknG exhibits unconventional E1 with isopeptidase activities *in vitro*. (*A*) *In vitro* ubiquitin conjugation assay of UbcH7. UbcH7 was incubated with GST-tagged PknG, Ub, and ATP/Mg^2+^ at 37 °C for 30 min. Reaction products were immunoblotted with antibodies against UbcH7 (top panel), Ub (middle panel) and the GST tag on PknG (bottom panel). (*B*) *In vitro* ubiquitin conjugation assay of UbcH7 catalyzed by PknG or its mutants. Reaction products were immunoblotted as in (*A*). (*C*) *In vitro* ubiquitin conjugation assay of UbcH7 catalyzed by PknG or its E1 activity-dead mutant (PknG P57A). Reaction products were immunoblotted as in (*A*). (*D*) MS/MS analysis of UbcH7 in the absence (top) or presence (bottom) of PknG to reveal ubiquitination sites. Tryptic peptide spanning residues ranging from 74 to 100 were identified as the high confidence peptide with di-Gly modification. The ubiquitinated residue (Lys82) in the peptide sequence is highlighted in red. The peak heights are the relative abundances of the corresponding fragmentation ions. Matched amino terminus-containing ions (b ions) are indicated in red, Matched carboxyl terminus-containing ions (y ions) are indicated in blue. Only the major identified peaks are labeled. (*E*) *In vitro* ubiquitin discharge assay of UbcH7∼Ub with or without PknG. Reaction products were immunoblotted as in (*A*). (*F*) *In vitro* ubiquitin conjugation assay of UbcH7 by PknG, PknG P57A, or PknG I87A. Reaction products were immunoblotted as in (*A*). (*G*) *In vitro* ubiquitin discharge assay of UbcH7∼Ub by PknG or its isopeptidase activity-dead mutant (PknG I87A). Reaction products were immunoblotted as in (*A*).

UbcH7∼Ub charged by canonical E1 (UbcH7 C86-Ub) can transfer Ub to free Cys, but not Lys, independent of an E3 (29, 30). We then further incubated UbcH7∼Ub charged by PknG (UbcH7 K82-Ub) with free Lys or Cys, and found that UbcH7 K82-Ub cannot directly transfer Ub to either free Lys or free Cys *in vitro* (*SI Appendix*, Fig. S3*C*). Next, we showed that PknG could also promote the discharge of Ub from UbcH7 K82-Ub (Fig. 2*E*). Because K82-Ub is linked by an isopeptide bond, this discharge was probably caused by an isopeptidase activity of PknG. Indeed, the isopeptidase activity of PknG could be greatly reduced by ubiquitin-aldehyde (Ub-Ad), a general inhibitor of isopeptidases (*SI Appendix*, Fig. S3*D*). We thus tried to identify the key sites for the isopeptidase activity of PknG. Among these PknG mutations, the PknG Ile87Ala (PknG I87A) mutation led to the accumulation of a larger quantity of UbcH7 Lys82-Ub, suggesting that Ile87 is required for the isopeptidase activity of PknG (Fig. 2*F* and *SI Appendix*, Fig. S3*E* and *F*). We then repeated the Ub discharge assay *in vitro*, and confirmed that PknG I87A mutation indeed abolished the isopeptidase activity of PknG (Fig. 2*G*). Together, Mtb PknG possesses unconventional E1 activity as well as isopeptidase activity.

### Mtb PknG inhibits host innate immunity by preventing NF-κB activatio

Ubiquitination system plays an important role in host innate immune response regulation, and thus emerges as a prime eukaryotic host target for bacterial pathogens (31, 32). We then sought to investigate whether Mtb PknG modulates host innate immune signaling pathways including NF-κB or MAPK pathways. Luciferase reporter assay was performed in human HEK293T cells transfected with vector encoding different mutants of PknG as well as fluorescent protein-reporting vectors. Transient expression of Mtb PknG in HEK293T cells markedly suppressed TNF-induced NF-κB pathway activation, while it had a slight effect on RacL61-induced AP-1 (a transcription factor of JNK and p38 MAPK pathways) activation and no effect on RasV12-induced Erk pathway activation (*SI Appendix*, Fig. S4 *A–C*). Notably, the E190A mutation, which disrupted the interactions between PknG and UbcH7, abolished the NF-κB activation-suppressive effects of PknG, while other PknG mutations including K181M and PknG ΔTPR did not have such effects (Fig. 3*A*). To better elucidate the specific regulatory function of PknG, we next constructed multiple PknG mutants including *ΔpknG:pknG K181M, ΔpknG:pknG E190A* and *ΔpknG:pknG ΔTPR*. The results demonstrated that all those PknG mutant Mtb strains grew at similar rates as the WT Mtb strain in 7H9 medium, and the PknG protein levels secreted by those mutants were also comparable to those from the WT Mtb strain (*SI Appendix*, Fig. S4 *D* and *E*). We then performed quantitative real-time PCR and enzyme-linked immunosorbent assay (ELISA) to examine whether the expression of cytokines was regulated by Mtb PknG or its mutants during Mtb infection. Deletion of PknG in Mtb promoted the production of inflammatory cytokines including TNF and IL-6 and decreased bacterial survival in primary human monocyte-derived macrophage cells, suggesting that PknG inhibits inflammatory cytokine production and enhances bacterial intracellular survival. The E190A mutation, but not the K181M mutation, largely abolished Mtb PknG-mediated suppression of inflammatory cytokine production in macrophage cells, indicating that PknG inhibits inflammatory cytokine production in Mtb-infected macrophage cells depending on its Ubl domain, but not its kinase activity (Fig. 3 *B* and *C*, and *SI Appendix*, Fig. S5). We also noticed that the E190A mutation largely abrogated, while the K181M mutation partially abrogated, the Mtb PknG-mediated promotion of bacterial survival in macrophage cells, indicating that PknG promotes the intracellular survival of Mtb mainly depending on its Ubl domain and partially depending on its kinase activity, the latter of which might regulate other cellular functions such as pathogen clearance rather than regulating the NF-κB pathway (Fig. 3*D*). Taken together, Mtb PknG could specifically inhibit NF-κB activation in a manner depending on its Ubl domain.

**Fig. 3.**
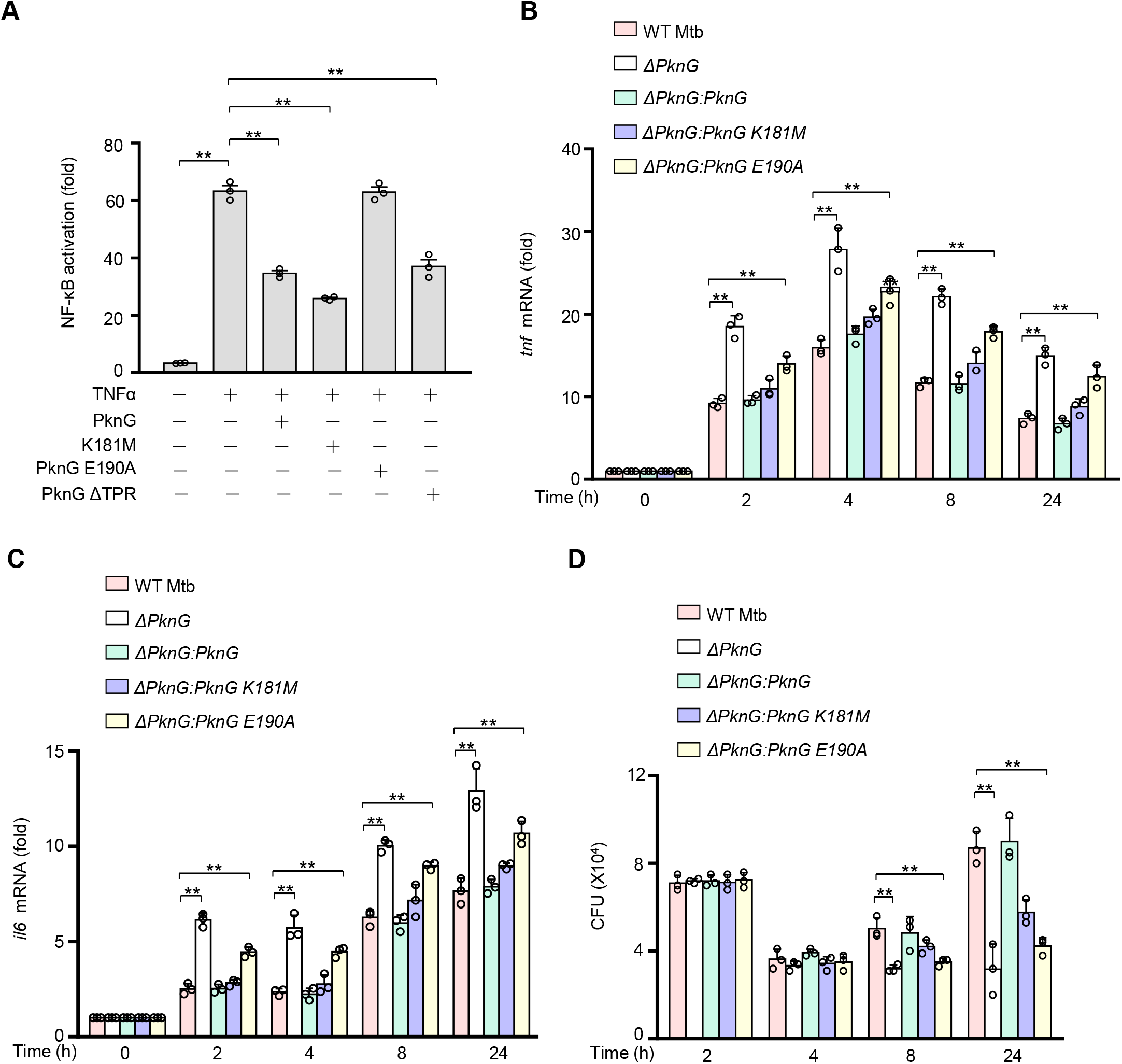
Mtb PknG inhibits NF-κB activation in a Ubl domain-dependent manner. (*A*) Luciferase assay of NF-κB activation in HEK293T cells transfected with WT or mutant Mtb PknG. (*B* and *C*) Quantitative PCR analysis of *Tnf* mRNA (*B*) and *Il6* mRNA (*C*) in primary human monocyte-derived macrophages. Cells were infected with WT, *ΔpknG, ΔpknG:pknG, ΔpknG:pknG K181M*, or *ΔpknG:pknG E190A* Mtb strain at an MOI of 1 for 0–24 h. The fold changes obtained from each sample was normalized to that of *GAPDH*. (*D*) Survival of Mtb strains in primary human monocyte-derived macrophages treated as in (*B*) and (*C*). Data are shown as mean ± SEM of at least three replicates. \**P* < 0.05 and ** *P* < 0.01 (one-way ANOVA). CFU, Colony-Forming units.

### Mtb PknG promotes the degradation of TRAF2 and TAK1 by serving as both E1 and E3

To explore the molecular mechanism by which PknG inhibits NF-κB signaling activation, we conducted further luciferase reporter assays in HEK293T cells and found that PknG suppressed the activation of NF-κB mediated by TAK1 and TAB1 (Fig. 4*A*). Co-immunoprecipitation analysis both in transfected HEK293T cells and infected macrophages further demonstrated that PknG interacted with TRAF2 and TAK1, but not TAB2 and TAB3, all of which are key components of the NF-κB signaling (33), through its UbcH7-binding domain (Fig. 4 *B* and *C* and *SI Appendix*, Fig. S6*A*). Consistently, pull-down assays also showed that TRAF2 and TAK1 bound to the Ubl domain of PknG both in the absence and presence of UbcH7 (*SI Appendix*, Fig. S6 *B–E*). Meanwhile, immunoblotting analyses indicated that the protein levels of TRAF2 and TAK1 markedly decreased in U937 cells infected with WT Mtb or Mtb *ΔpknG:pknG* strain compared with those infected with Mtb *ΔpknG* or Mtb *ΔpknG:pknG E190A* strain (Fig. 4*D*), suggesting that PknG might promote protein degradation of TRAF2 and TAK1 through its unconventional ubiquitinating activity. Accordingly, PknG and its K181M mutant, but not its E190A mutant, promoted UbcH7-dependent polyubiquitination of TRAF2 and TAK1 in infected macrophages (Fig. 5 *A* and *B*). Similar results were obtained in transiently transfected HEK293T cells (*SI Appendix*, Fig. S6 *F* and *G*). Furthermore, the addition of UbcH7 and Ub to the purified TRAF2 or TAK1 promoted the ubiquitination of TRAF2 or TAK1 *in vitro* in the presence of WT PknG, but not its E1 or isopeptidase activity-dead mutant (Fig. 5 *C* and *D*). We then showed that PknG promoted the K48-linked, but not K63-linked, polyubiquitination of TRAF2 and TAK1 (Fig. 5 *E* and *F*). Taken together, Mtb PknG processes unconventional E1 and E3 activities to promote ubiquitination of TRAF2 and TAK1, leading to their ubiquitination-mediated degradation and the ensuing inhibition of the NF-κB pathway.

**Fig. 4.**
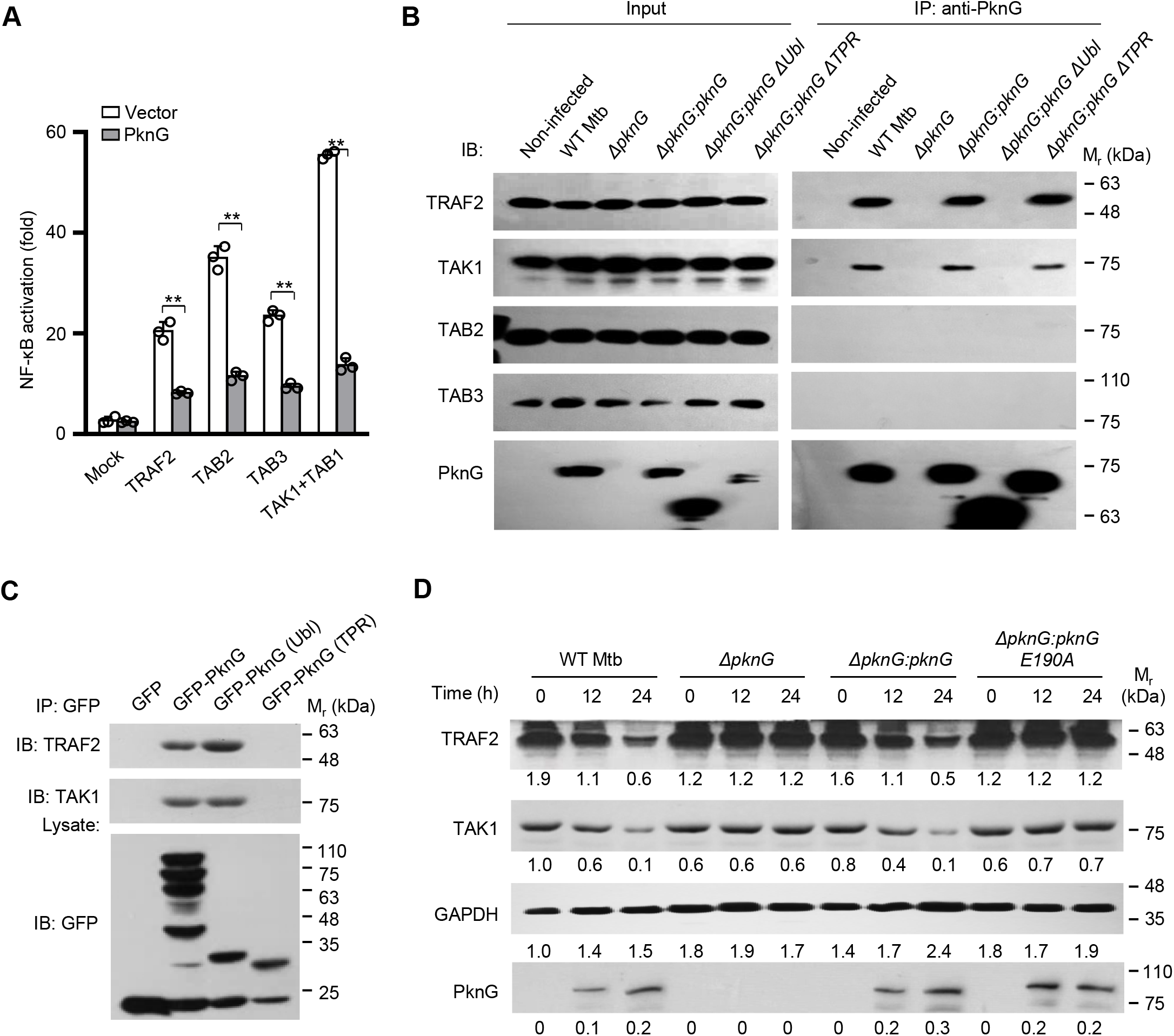
Mtb PknG targets TRAF2 and TAK1 for degradation by binding to UbcH7. (*A*) Dual-luciferase assay of TRAF2-, TAB2-, TAB3-, TAK1-, or TAB1-induced NF-κB activation in the absence or presence of Mtb PknG in HEK293T cells. (*B*) IP of TRAF2 and TAK1 by Mtb PknG in U937 cells. Cells were infected with the indicated Mtb strains at an MOI of 1. Non-infected cells were used as control. After 4 h, cells were lysed and immunoprecipitated with antibody against PknG. The immunoprecipitated proteins were immunoblotted with TRAF2, TAK1, TAB2 (negative control), or TAB3 (negative control) antibody. (*C*) IP of TRAF2 and TAK1 by GFP-tagged Mtb PknG or its truncated forms in HEK293T cells. (*D*) Immunoblotting analysis of TRAF2 and TAK1 from lysates of U937 cells infected with the indicted Mtb strains as in (*B*) for 0–24 h. Densitometry quantifications of immunoblots are indicated below the immunoblots. Data are shown as mean ± SEM of three replicates in (*A*). \**P* < 0.05 and ** *P* < 0.01 (two-tailed unpaired *t*-test).

**Fig. 5.**
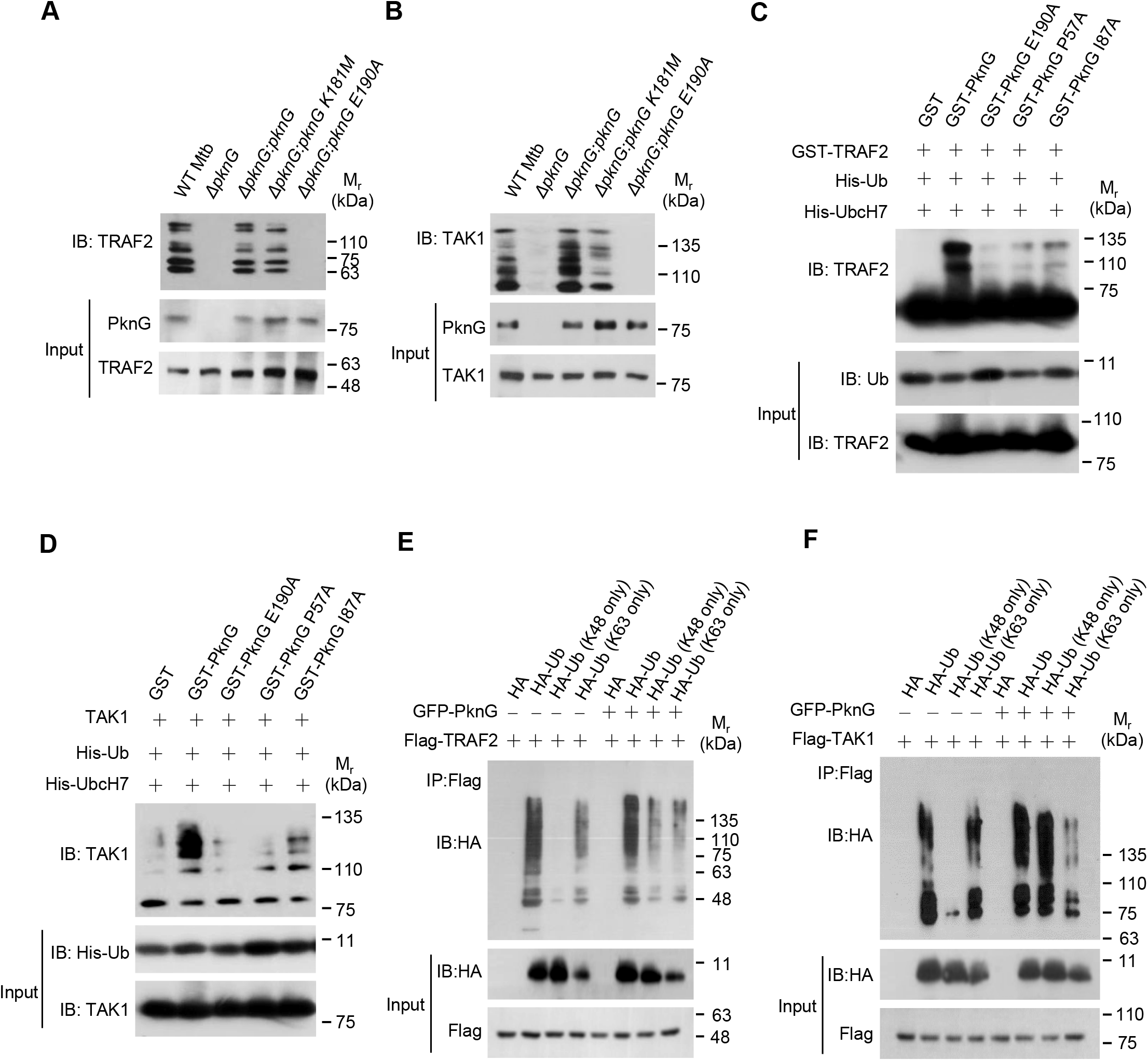
Mtb PknG promotes Lys48-linked poly-ubiquitination of TRAF2 and TAK1 depending on UbcH7-binding. (*A* and *B*) *In vivo* ubiquitination assay of TRAF2 (*A*) and TAK1 (*B*) in U937 cells infected with WT, *ΔpknG, ΔpknG:pknG, ΔpknG:pknG K181M*, or *ΔpknG:pknG E190A* Mtb strains at an MOI of 1 for 6 h. Endogenous ubiquitinated proteins were enriched using TUBE reagents and immunoblotted with antibodies against TRAF2 (*A*, top panel) or TAK1 (*B*, top panel). (*C* and *D*) *In vitro* ubiquitination of TRAF2 (*C*) and TAK1 (*D*) in the presence of PknG or its mutants. Reaction products were immunoblotted with antibodies against TRAF2 (*C*, top panel) or TAK1 (*D*, top panel). (*E* and *F*) *In vivo* ubiquitination assay of TRAF2 (*E*) and TAK1 (*F*) in the absence or presence of PknG in HEK293T cells transfected with the indicated plasmids for 24 h. cells were lysed and immunoprecipitated with antibody against Flag. The immunoprecipitated proteins were immunoblotted with antibody against HA.

### Mtb PknG suppresses immune responses to mycobacteria *in vivo*

To further examine the contribution of the unconventional E1 and E3 activities of PknG in host innate immune suppression during mycobacterial infection *in vivo*, we challenged C57BL/6 mice intratracheally with WT Mtb and different PknG mutant Mtb strains. Lungs, livers and spleens were harvested and analyzed at day 0, 5, 10, 15 and 20 after infection. Quantitative real-time PCR of splenic cells revealed that the *ΔpknG:pknG E190A* strain markedly increased the levels of *Tnf* and *Il6* mRNAs in spleens of the infected mice compared with the WT Mtb strain (Fig. 6 *A* and *B*). CFU counting and hematoxylin and eosin staining were then performed to evaluate pathologic changes in lungs or livers. Consistently, the *ΔpknG:pknG E190A* strain decreased bacterial load in lungs and reduced cellular infiltration in lungs and livers of the infected mice at day 10 after infection as compared to WT Mtb strain (Fig. 6 *C* and *D*). Collectively, the UbcH7-binding-dependent E1 and E3 enzyme activities of PknG contribute to its innate immune suppression function *in vivo*.

**Fig. 6.**
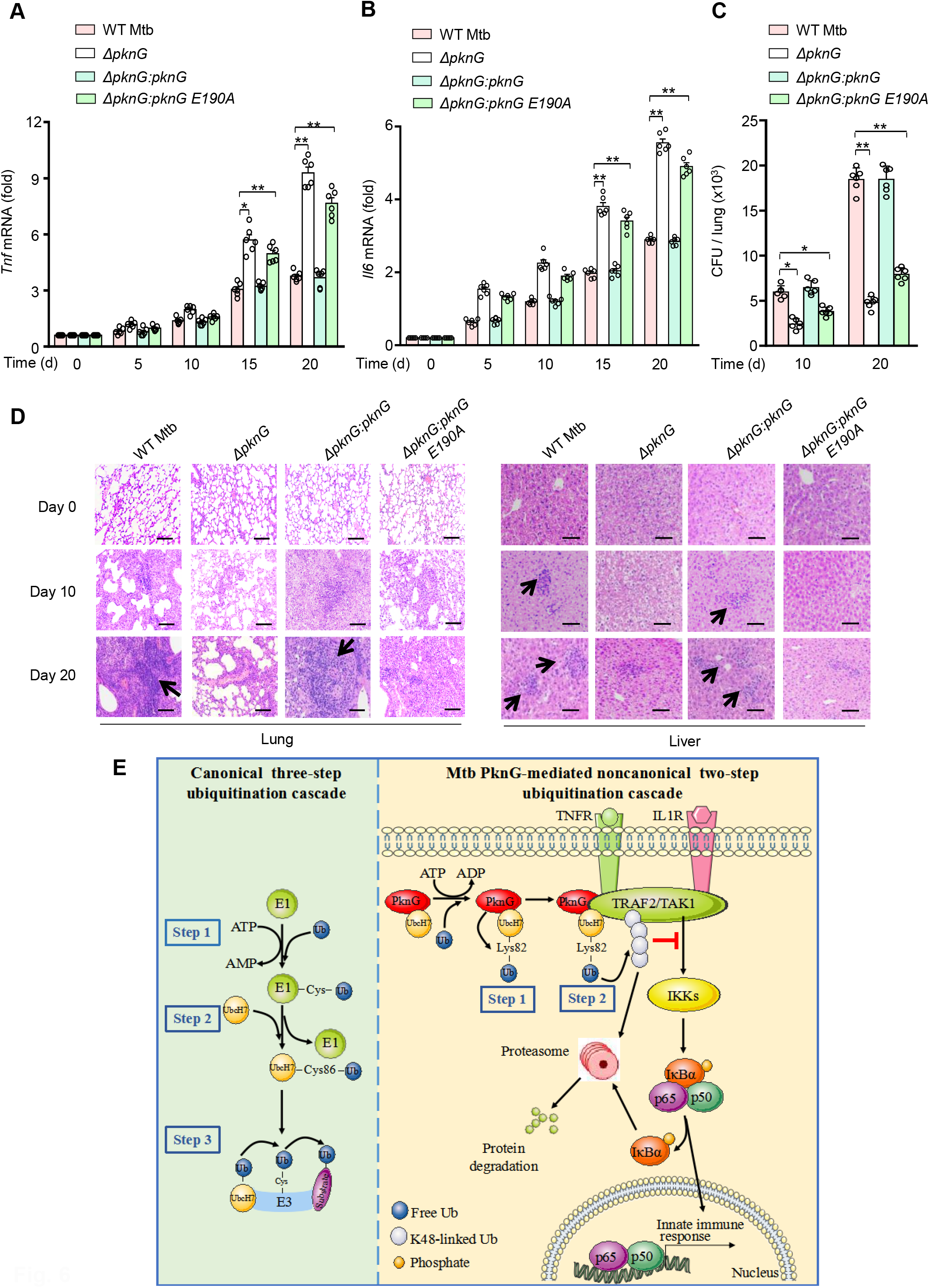
The Ubl domain of PknG contributes to host innate immune suppression i*n vivo*. (*A* and *B*) Quantitative PCR analysis of *Tnf* mRNA (*A*), *Il6* mRNA (*B*) in splenic cells from C57BL/6 mice infected with WT or mutant Mtb stains for 0–20 days. (*C*) Bacterial load in lungs from C57BL/6 mice treated as in (*A*). (*D*) H&E staining of lungs and livers of C57BL/6 mice treated as in (*A*). Arrows indicate puncta of cellular infiltration. Scale bars, 200 μm. (*E*) Proposed model depicting Mtb PknG-mediated host innate immune suppression. During mycobacterial infection, PknG binds ATP to catalyze a unique two-step ubiquitination cascade through unconventional E1 and E3 enzyme activities with the release of ADP instead of AMP, leading to degradation of TRAF2 and TAK1 and suppression of host innate immune responses. Data are shown as mean ± SEM of three replicates in (*A*) to (*C*). \**P* < 0.05 and ** *P* < 0.01 (one-way ANOVA).

## Discussion

Mtb encounters a hostile environment within host cells during the course of infection. As the most successful intracellular bacterial pathogen, Mtb not only has the ability to adapt to a changing host environment, but also actively interferes with various host signaling pathways and cellular functions to counteract, inhibit, or hijack various killing apparatus employed by host cells (2, 34-36). The intimate and extensive association of Mtb with its host seems to be the driving force in the evolution of its pathogenicity and offers a fascinating peek into how pathogens exploits host cellular functions to their advantage (34). One type of powerful weapons developed by Mtb for sensing and interacting with its host are a set of eukaryotic-type serine/threonine protein kinases and protein tyrosine phosphatases that are potential targets for TB treatment (5, 31, 37, 38). It was shown that Mtb PknG could block phagosome maturation and pathogen clearance in a kinase activity-dependent manner (3, 8). In this study, adopting a multidisciplinary approach containing multiple molecular and cellular methodologies as well as mice infection models, we revealed additional important host regulatory roles of Mtb PknG in suppressing innate immune signaling. More strikingly, we found that the eukaryotic-type kinase PknG exhibits additional unconventional E1 and E3 enzyme activities, which initiate host regulatory function on innate immune responses during Mtb infection. Thus, PknG represents a powerful mycobacterial effector protein that could exert multifaceted regulatory roles in host cellular functions including innate immune signaling and phagosome maturation, depending on its distinct eukaryotic-like enzymatic activities conferred by its specific eukaryotic-like domains. Such highly flexible and multipronged host-regulatory functions of the pathogen effector protein might help the pathogen better adapt to and survive within the highly changeable and complicated intracellular environment (39). The intricate tactics adopted by those pathogen effector proteins could also teach us new knowledge of cell biology since they act as natural inhibitors or agonists that target host signaling pathways and cellular functions in an unusual but normally highly efficient manner.

One of the strategies adopted by the pathogenic bacteria to promote their intracellular survival is to secrete diversified effector proteins into host cells. Once entered into the host cells, those effectors could subvert various cellular processes to destroy host immune defenses. Notably, the ubiquitination pathway is among one of the major targets of many bacterial effectors, since ubiquitination is indispensable for eukaryotic cells to regulate diverse immune responses. In addition to regulating the function of crucial enzymes (such as host E1s, E2s, E3s, and DUBs) involved in host ubiquitination pathway, bacterial pathogens have also evolved certain eukaryotic-like enzymes to regulate ubiquitination process (19, 40). Previous studies suggested that charged E1∼Ub recruits UbcH7 and transfers Ub to the conserved Cys86 residue of UbcH7 (UbcH7 C86-Ub), which then interacts with HECT (homologous with E6-associated protein C-terminus)- and RBR (RING-Between-RING)-type E3s to form an intermediate E3∼Ub thiol-ester bond on the conserved cysteine residues in those E3s (41). Here, we surprisingly found that Mtb PknG possesses both E1 and E3 enzymatic activities towards host substrate proteins, including TRAF2 and TAK1, during Mtb infection. Furthermore, unlike any of those previously reported E1s and E3s, Mtb PknG exerts dual-functional E1 and E3 activities in a unique and sequential way. Specifically, PknG initially binds to the E2 protein UbcH7 via a novel Ubl domain and exerts unconventional E1 activity to directly catalyze Ub conjugation to the Lys82 site of UbcH7 (UbcH7 K82-Ub), instead of forming UbcH7 C86-Ub thiol-ester bond. PknG further acts as an isopeptidase to promote the discharge of Ub from UbcH7 K82-Ub, and then transfers Ub to the recruited substrates (including TRAF2 and TAK1), leading to ubiquitination-mediated degradation of TRAF2 and TAK1 and the ensuing suppression of NF-κB signaling. A typical E3 usually binds its E2 and substrates using distinct domains or regions. Notably, the Ubl domain of PknG binds its substrates in addition to its E2 UbcH7, suggesting that the Ubl domain of PknG possesses an unidentified substrate-binding region. Thus, PknG represents the first ubiquitinating enzyme to promote the attachment of Ub to an E2 via an isopeptide bond by acting as both E1 and E3. More strikingly, PknG binds ATP to catalyze E2-Ub conjugation via an isopeptide bond with the release of ADP instead of AMP, a phenomenon different from the conventional E1 that usually hydrolyzes ATP to release AMP (25-28). A better understanding of the intimate cross-talk, antagonism and co-evolution of the Mtb and its host could be exploited in the development of novel TB treatments. Such pathogen-host interfaces-based novel therapies could be effective for both drug-susceptible and drug-resistant TB, and such strategy of targeting secreted pathogen effectors instead of pathogen essential genes could help minimize further development of mycobacterial drug resistance, which is important against the backdrop of growing challenges of antibacterial drug resistance worldwide (42, 43).

In summary, our study demonstrates that Mtb PknG possesses unconventional E1 and E3 activities, attacking host innate immunity by catalyzing a novel two-step ubiquitination cascade (Fig 6*E*). Importantly, we found that the Ubl domain of Mtb PknG was highly conserved among Mtb clinical isolates as analyzed using the GMTV database (http://mtb.dobzhanskycenter.org) (Dataset S2) (44). We also noticed that the Ubl domain (especially its E190 site) of Mtb PknG exhibited high homology to some other pathogenic mycobacteria, but little homology to non-pathogenic mycobacteria and no homology to human hosts as analyzed against the Uniport Reference Proteomes database (Dataset S3). This feature might make PknG selectively targeted. The findings from this study might suggest a potential TB treatment via targeting the unconventional E1 and E3 activities of Mtb PknG.

## Materials and Methods

### Bacterial strains, mammalian cell lines and plasmids

*E. coli* DH5α and BL21 (DE3) were used for genetic manipulation and protein overexpression. *Mycobacterium tuberculosis* (Mtb) H37Rv was used for infection. PknG deletion and mutant strains in Mtb were created as described previously (32). Mtb strains included wild-type (WT) Mtb, *pknG*-deleted Mtb (*ΔpknG*), Mtb *ΔpknG* complemented with WT *pknG* (*ΔpknG:pknG*), *pknG K181M* (*ΔpknG:pknG K181M*, which is a kinase-dead mutant), *pknG E190A* (*ΔpknG:pknG E190A*, which is a mutant that does not interact with UbcH7), and *pknG ΔTPR (ΔpknG:pknG ΔTPR*, from which the TPR domain was deleted). Gene sequencing was used to confirm the mutation, deletion or truncation of PknG in Mtb strains. Expression of PknG in infected host cells was examined by immunoblotting analysis. HEK293T (ATCC CRL-3216) and the human monocytic cell line U937 cells (ATCC CRL-1593.2) were obtained from the American Type Culture Collection (ATCC). For expression in mammalian cells, *pknG* or its mutants were amplified by PCR and cloned into pEGFP-N1 (with GFP-tag) or p3xFlag-CMV14 (with Flag-tag) vectors. Bacterial expression plasmids were constructed by cloning the cDNA into pET30a (with His_6_-tag) or pGEX-6P-1 (with GST-tag). Site-directed mutagenesis of *pknG* was performed using the Mut Express II Fast Mutagenesis Kit V2 (Vazyme). All the plasmids were sequenced at the Beijing Genomics Institute (BGI) for verification. The strains, plasmids and oligonucleotides used in this study are listed in Table S1.

### Antibodies

All of the antibodies were used according to manufacturer’s instructions and based on previous experiences in the lab. Rabbit anti-PknG antibody was produced and purified by GenScript Biotechnology with the recombinant GST-tagged PknG protein as immunogen (1/10000 for immunoblotting, 1/200 for Immunohistochemistry). The following commercial antibodies were used: anti-UbcH7 (A300-737A, Bethyl), 1:2000 for immunoblotting; anti-Flag (F1804, Sigma), 1:5000 for immunoblotting; anti-GST (TA-03, ZSGB-BIO), 1:5000 for immunoblotting; anti-ubiquitin (131600, Invitrogen), 1:2000 for immunoblotting; anti-His (TA-02, ZSGB-BIO), 1:2000 for immunoblotting; anti-GFP (ab1218, Abcam), 1:2000 for immunoblotting; anti-TRAF2 (SAB3500375, Sigma),1:2000 for immunoblotting; anti-TAK1 (12330-2-AP, Proteintech),1:1000 for immunoblotting; anti-GAPDH (sc-25778, Santa Cruz), 1:5000 for immunoblotting; anti-HA (3724, Cell signaling), 1:2000 for immunoblotting; anti-Rv3134 (sc-52107, Santa Cruz), 1:1000 for immunoblotting; anti-Ag85B (ab43019, Abcam), 1:1000 for immunoblotting; anti-TAB3 (sc-67112, Santa Cruz Biotechnology), 1:1000 for immunoblotting; anti-TAB2 (BF 0376, Affinity Biosciences), 1:1000 for immunoblotting.

### Expression and purification of recombinant proteins

For GST or His fusion proteins, *E. coli* BL21 (DE3) strain harboring pGEX-6P-1 or pET30a derivatives were cultivated in LB medium supplemented with antibiotic for 3 h at 30 °C. Protein expression was induced by 1 mM IPTG for 3 h at 30 °C. GST fusion protein and His fusion protein were purified with glutathione-Sepharose 4B (GE Healthcare) and Ni-NTA Agarose (QIAGEN) separately according to the manufacturer’s protocol. Protein concentrations were determined by the BCA protein assay (Pierce; Rockford) and protein purity was examined by SDS-PAGE analysis.

### Cell culture and bacteria cultivatio

HEK293T cells were cultured in Dulbecco’s modified Eagle medium (Hyclone) supplemented with 10% fetal calf serum (FBS). U937 cells were maintained in RMPI 1640 medium with 10% FBS. Before infection, U937 cells were differentiated into adherent macrophage-like cells using 10 ng/ml PMA (Sigma) for 24 h. *E. coli* DH5α and BL21 (DE3) were grown in LB medium. Mtb strains were cultivated in middlebrook 7H9 broth supplemented with 10% oleic acid-albumin-dextrose-catalase (OADC) and 0.05% Tween-80 (Sigma), or on Middlebrook 7H10 agar (BD) supplemented with 10% OADC.

### Preparation of U937 cells and primary human monocyte-derived macrophage cells

U937 cells were differentiated with 10 ng/ml PMA and washed twice with PBS, and further cultured in fresh medium for an additional 2 h prior to infection. Peripheral blood mononuclear cells were prepared from venous blood from healthy volunteer donors using density gradient centrifugation. All experiments involving human subjects were approved by the Biomedical Research Ethics Committee of the Institute of Microbiology, Chinese Academy of Sciences. Written informed consent was obtained from all donors. For differentiation into macrophages, monocytes (2 × 10^6^) were cultured in 500 µl RPMI1640 with 10% FBS, 4 mM L-glutamine and 1% penicillin-streptomycin (Invitrogen) in 24-well plates for 7 days. Recombinant human macrophage colony-stimulating factor (M-CSF) was added at day 0, 2, and 4. At day 7, macrophages were used for mycobacterial infection.

### Quantitative real-time PCR

Primary human monocyte-derived macrophages were infected with WT, *ΔpknG, ΔpknG:pknG, ΔpknG:pknG K181M*, or *ΔpknG:pknG E190A* Mtb strain at an MOI of 1 for 0–24 h. Total RNA was extracted from the infected U937 cells using Total mRNA isolation kit (R1061, Dongsheng), and was reverse-transcribed into cDNA using the Hieff First Strand cDNA Synthesis Super Mix (11103ES70, YEASEN). The cDNA was then analyzed by qRT-PCR using Hieff qPCR SYBR Green Master Mix (11202ES08, YEASEN) on ABI 7500 system (Applied Biosystems). Each experiment was performed in triplicates and repeated at least three times. Data were analyzed by the 2^−ΔΔCT^ method and were normalized to the expression of the control gene *GAPDH*.

### Ub-charging reactio

Reaction mixtures containing 2 µM GST-tagged PknG or its mutants (purified from *E. coli*), 5 µM Ub (U-100H, Boston Biochem), 5 µM UbcH7 (E2-640, BostonBiochem) in reaction buffer [50 mM Tris/HCl pH 7.4, 10 mM MgCl_2_, 2 mM ATP (A6559, Sigma) or ATP analogs including APCPP (M6517, Sigma), AMPPNP (A2647, Sigma), and ATP-γ-S (A1388, Sigma)], were incubated at 37 °C for 30 min. Samples were quenched in non-reducing loading buffer and analyzed by SDS–PAGE.

### Mass spectrometry for determination of ubiquitination sites

Bands were excised from the SDS-PAGE gel, rinsed with 100 mM ammonium bicarbonate, dried and reduced using 10 mM dithiothreitol (DTT) in 100 mM ammonium bicarbonate at 56 °C, and alkylated with 55 mM iodoacetamide. Proteins were then digested with 12.5 ng/ul trypsin overnight at room temperature. After trypsinization, the proteins were eluted from the gel by alternating four times between 50% acetonitrile, 5% formic acid, and 50 mM sodium bicarbonate and an additional three times alternating between 100% acetonitrile and 50 mM NH_4_CO_3_. Samples were then desalted on a C18 ZipTip pipette tip (Millipore) and lyophilized for mass spectrometry (MS) analysis. Peptides were separated via Nano LC-nano MS/MS analysis on a NanoAcquity system (Waters, Milford, MA) connected to an LTQ-Orbitrap XL hybrid mass spectrometer (Thermo Fisher Scientific, Bremen, Germany). The MS and MS/MS raw data were processed by using Proteome Discoverer (Version 1.4.0.288, Thermo Fischer Scientific) and searched against the customized Uniprot_*M. tuberculosis*_GNCC_2019 database. Peptide ubiquitination sites were identified via protein database searching of the resulting tandem mass spectra using Mascot. The Mascot search engine was used to identify ubiquitinated residues marked as LeuArgGlyGly (LRGG) or GlyGly (GG). The mass spectrometry data have been deposited in the Proteomics Identification Database (PRIDE) under an accession numbers PXD020982.

### Nucleophile reactivity assays

Reactions containing 2 µM GST-tagged PknG or E1 (purified from *E. coli*), 5 µM Ub (U-100H, Boston Biochem) and 5 µM UbcH7 (E2-640, BostonBiochem) in reaction buffer (50 mM Tris/HCl pH 7.4, 10 mM MgCl_2_/ATP) were incubated at 37 °C. After 30 min, GST-E1 and GST-PknG were removed from the reactions with glutathione-Sepharose 4B (GE Healthcare). Next, Reaction mixtures containing 5 µM ubiquitin conjugation UbcH7 (UbcH7∼Ub) (charged by GST-tagged E1 or Mtb PknG) and 20 mM of L-lysine monohydrochloride in 50 mM Tris/HCl buffer (pH 7.4) in the presence or absence of PknG or PknG I87A mutant were incubated for 0–60 min at 37 °C. Samples were quenched in non-reducing loading buffer and analyzed by SDS–PAGE.

### *In vitro* isopeptidase activity assay

Reaction mixtures containing 1 µM of ubiquitin-conjugated UbcH7 (UbcH7∼Ub) (which is charged by GST-tagged Mtb PknG), 2 nM of purified GST or GST-PknG, and 20 mM of L-lysine monohydrochloride in 50 mM Tris/HCl buffer (pH 7.4) were incubated in the presence or absence of ubiquitin-aldehyde (6 nM, Abcam) for 30 min at 37 °C. Samples were resolved by 10% SDS-PAGE.

### Luciferase reporter assay

Luciferase reporter assay was performed in the presence or absence of 1 μg of PknG plasmid as described previously (32). The fold induction was calculated as [relative FU stimulated]/ [relative FU unstimulated].

### Pull-down assay

The recombinant proteins (20 μg) were added to 20 μl of glutathione resin (for GST-tagged proteins) in 500 μl of binding buffer (50 mM Tris, pH 7.5, 150 mM NaCl, 5 mM DTT and 0.1% NP-40) supplemented with 1% protease inhibitors for 1 h at 4°C. The beads were washed five times with binding buffer and further incubated with prey protein supplemented with 0.1 mg/ml BSA (the molar ratio of GST-tagged protein and prey protein is 1:1). After 2 h of incubation at 4 °C, beads were washed and subjected to immunoblotting.

### Immunoblotting, CFU counting, quantitative PCR, and enzyme-linked immunosorbent assay (ELISA) of infected macrophage cells

Macrophage cells were infected with Mtb strains at a multiplicity of infection (MOI) of 1 for 0–24 h. Immunoblotting, CFU counting, quantitative PCR and ELISA of infected macrophage cells were performed as described previously (32). PCR primers used are listed in Table S1. ELISA Kits (Human TNF ELISA kit: ELH-TNF-1, Ray biotech; Human IL-6 ELISA kit: ELH-IL6-1, Ray biotech) were used for ELISA analysis according to the manufacturer’s instructions.

### Immunoprecipitatio

For assays with HEK293T, cells were transfected using Lipo 2000 (Invitrogen) for 24 h. For assays with U937, cells were infected with Mtb strains at an MOI of 1. After 1 h, cells were washed with 1× PBS for three times to exclude non-internalized bacteria, and were then incubated in the fresh RPMI 1640 medium for additional 4 h. Both HEK293T and U937 cells were lysed in Western and Immunoprecipitation (IP) buffer (Beyotime) for 30 min at 4 °C. Cell lysates were then incubated with Flag M2 beads (Sigma), anti-HA affinity matrix (Roche), GFP-Nanoab-Agarose (V-Nanoab), or PknG antibody-protein A/G agarose at 4 °C, followed by extensive wash with the Western and IP buffer. The immunoprecipitated samples were analyzed by immunoblotting with indicated antibodies.

### Yeast two-hybrid assay

Yeast two-hybrid assay was performed using the Matchmaker Two-Hybrid System (Clontech) by following the manufacturer’s instructions. A mouse 7-day embryo cDNA Library (CATALOG No. 630478; Clontech Laboratories, Inc.) was used to identify host interaction proteins of Mtb PknG through yeast two-hybrid assay. Mtb PknG gene was subcloned into the plasmid pGBKT7 as the bait plasmid. *Saccharomyces cerevisiae* AH109 cells were cotransduced with the bait plasmids and the prey plasmids by the lithium acetate method. To test the interactions between proteins, the transformants were streaked onto low-stringency (lacking leucine and tryptophan) and high-stringency (lacking adenine, histidine, leucine and tryptophan) selection plates.

### *In vitro* ubiquitination assays

*In vitro* ubiquitination assays of GST-tagged TRAF2 (ab204091, Abcam) and TAK1(ab132950, Abcam) were performed in 40 μl reaction mixture containing reaction buffer (25 mM Tris-HCl [PH 7.5], 50 mM NaCl and 10 mM ATP), 1 μg TRAF2 or TAK1, 2 μg UbcH7, and 2 μg His-Ub in the presence or absence of WT PknG or its mutants. Reactions were incubated at 30 °C for 2 h and reactions were terminated by adding SDS sample buffer without <u>β</u>-mercaptoethanol and heating at 100 °C for 5 min. Samples were resolved by SDS–PAGE on 10% gels and then subjected to immunoblotting.

### *In vivo* ubiquitination assays

*In vivo* ubiquitination assays were performed using U937 or HEK293T cells. For assays with U937, cells were infected with WT, *ΔpknG, ΔpknG:pknG*, or *ΔpknG:pknG E190A* Mtb strain at an MOI of 1 for 2 h. After another 2 h, the media were discarded. The cells were then washed three times with PBS to exclude non-internalized bacteria and cultured in fresh RPMI-1640 medium supplemented with 10 μM MG132 for an additional 4 h. Cells were lysed in Western and IP buffer (Beyotime) and centrifuged at 12,000*g*. Equilibrated Agarose-TUBEs (10–20 µl) were incubated with lysate at 4 °C. After 4 h, Agarose-TUBEs were washed for 5 times with TBS-T buffer and treated with 0.2 M glycine HCl (pH 2.5) for at least 1 h on a rocking platform at 4 °C. Agarose-TUBEs were then collected by high speed centrifugation (13,000*g*) for 5 min, and the supernatants were resolved by SDS–PAGE and then subjected to immunoblotting.

For assays with HEK293T, cells were transfected with Flag-TRAF2 (or Flag-TAK1) and other indicated plasmids for 24 h. Cells were lysed in Western and IP buffer (Beyotime) and centrifuged at 12,000g. The pre-washed anti-FLAG M2 beads were incubated with the supernatant for 2 h at 4 °C to pull-down Flag-tagged TRAF2 (or TAK1) and its conjugated proteins. Beads were then washed by Western and IP buffer and analyzed by immunoblotting.

### Murine infectio

C57BL/6 mice were purchased from Vital River (Beijing) and were kept using standard humane animal husbandry protocols. Mice were 6–8-week old during the course of the experiments and were age- and sex-matched in each experiment. Sample size was based on empirical data from pilot experiments. No additional randomization or blinding was used to allocate experimental groups. All experimental protocols were performed in accordance with the instructional guidelines of the China Council on Animal Care, and were approved by the Biomedical Research Ethics Committee of the Institute of Microbiology, Chinese Academy of Sciences, and the Beijing Chest Hospital, Capital Medical University. Mice were intratracheally infected with 1 × 10^5^ CFU of Mtb per mouse in 25 µl PBST. Lungs, livers, and spleens were harvested and analyzed as described previously (32). Staining with hematoxylin and eosin stain was performed for evaluation of tissue pathologic changes in a blinded fashion by a pathologist. The splenic cells were used for quantitative PCR.

### Data analysis

We analyzed Mtb *pknG* gene sequence conservation among Mtb clinical isolates using GMTV database (http://mtb.dobzhanskycenter.org). To acquire greater specificity and lower rate of false positive homologous proteins, we used phmmer tool based on hidden Markov model against Uniport Reference Proteomes database to predict homologous proteins of Mtb PknG in *homo sapiens* and all bacteria species. Only alignments with *E*-values less than 10^−2^ were considered homologous proteins.

### Statistics

Statistical analyses were performed using Prism 7.0 (GraphPad Software). All data were analyzed using an unpaired Student’s *t*-test or one-way ANOVA with Bonferroni post-test to correct for multiple compares as indicated in the corresponding figure legends. The results were averaged and expressed as means with SEM. Each value of \**P* < 0.05 or \*\**P* < 0.01 was considered to be statistically significant.

## Acknowledgements

We thank Xiang Ding (Laboratory of Proteomics, Core Facility of Protein Science, Institute of Biophysics, Chinese Academy of Sciences) and Ying Fu (Public Technology Service Center, Institute of Microbiology, Chinese Academy of Sciences) for the ubiquitination modification analysis with mass spectrometry. This work was supported by the National Natural Science Funds for Distinguished Young Scholar (81825014), the Strategic Priority Research Program of the Chinese Academy of Sciences (XDB29020000), the National Key Research and Development Program of China (2017YFA0505900 and 2017YFD0500300), the National Natural Science Foundation of China (31830003, 31530014, 81371769, 81871616 and 81571954), the National Science and Technology Major Project (2018ZX10101004), the Key Program of Logistics Research (BWS17J030).

## Author contributions

C.H.L. conceived and supervised the study. C.H.L., J.W., X.-B.Q. and G.F.G. designed the experiments. J.W., P.G. and Z.H.L. performed the majority of the experiments. Z.L., L.Q., Q.C., Y.Z., D.Z., B.L., J.S., Y.P. and Y.S. were involved in specific experiments. C.H.L., J.W. and X.-B.Q. analyzed the data and wrote the manuscript. All authors discussed the results and commented on the manuscript.

