## Supplementary materials for "A *Mycobacterium tuberculosis* effector protein attacks host innate immunity by acting as an unusual ubiquitinating enzyme"

**The PDF file includes:**

Supplementary text

Figures S1 to S6

Legends for Figures S1 to S6

Table S1

### Supplementary text

### Supplementary Materials and Methods

***In vitro* autophosphorylation of PknG and its mutants.** GST-tagged PknG or its mutants (5 μg) were incubated in 25 mM Tris (pH 7.5), 2 mM $MnCl_2$, 1 mM DTT and 200 μM ATP for 30 min at 37 ℃. The reactions were stopped by adding sample buffer and boiling for 10 min at 95 ℃. The samples were resolved by SDS–PAGE on 10% gels and then subjected to immunoblotting.

**LC-MS.** Reaction mixtures containing 2 μM GST-tagged PknG K181M (purified from *E. coli*), 5 μM ATP (A6559, Sigma) in reaction buffer [50 mM Tris/HCl pH 7.4, 10 mM $MgCl_2$], were incubated at 37 ℃ for 30 min in the presence or absence of 5 μM Ub (U-100H, Boston Biochem) and 5 μM UbcH7 (E2-640, BostonBiochem). Liquid chromatography-tandem mass spectrometry (LC-MS) data were acquired using a 6500 QTRAP triple quadrupole mass spectrometer (Applied Biosystems/Sciex) that was coupled to a ExionLC LC system (Applied Biosystems/Sciex) using Rezex RCM-Monosaccharide $Ca^+$ (Phenomenex, 300*7.8 mm). Compounds were eluted with isocratic elution of water with a flow rate of 0.4 mL/min over 40 min. MS analyses were carried out using electrospray ionization (ESI) and multiple reaction monitoring (MRM) scans in the negative ion mode. Injection volume was 5.00 μl. The dwell time for each transition was 50 ms, the ion spray voltage was 4.5 kV, and the source temperature was 550 ℃. File was created with the software Analyst (version 1.6.3, Applied Biosystems/Sciex). Peak area ratios of compounds were calculated using MultiQuant software (Version 3.0.2; Applied Biosystems/Sciex).

**Figure S1**

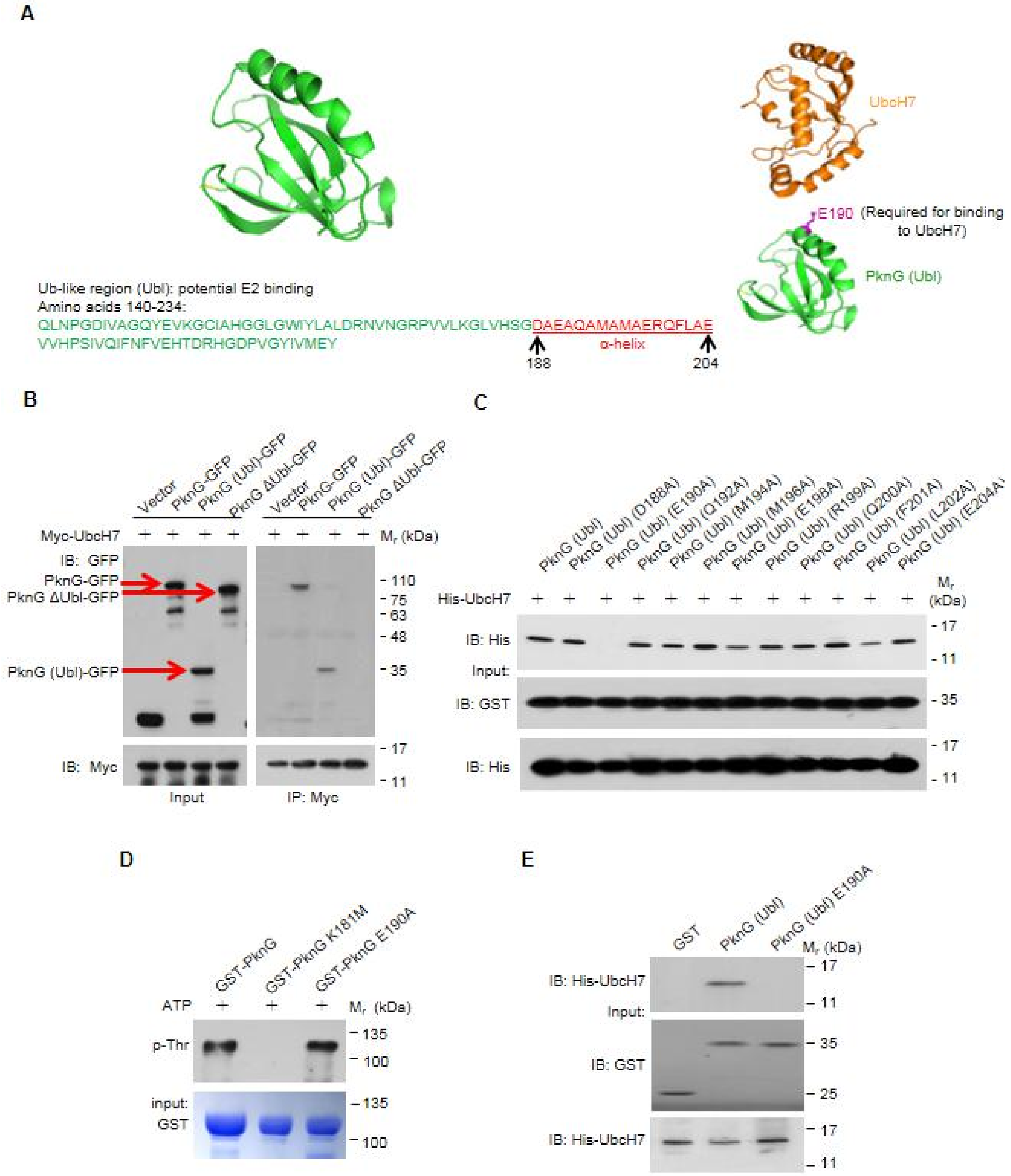
A
UbcH7
E190 (Required for binding to UbcH7)
PknG (Ubl)
Ub-like region (Ubl): potential E2 binding
Amino acids 140-234:
QLNPGDIVAGQYEVKGCIAHGGLGWIYLALDRNVNGRPVVLKGLVHSGDAEAQAMAMAERQFLAE
VVHPSIVQIFNFVEHTDRHGDPVGYIVMEY
α-helix
188
204
B
Vector
PknG-GFP
PknG (Ubl)-GFP
PknG ΔUbl-GFP
Myc-UbcH7
Mr (kDa)
IB: GFP
PknG-GFP
PknG ΔUbl-GFP
PknG (Ubl)-GFP
IB: Myc
Input
IP: Myc
110
75
63
48
35
17
11
C
PknG (Ubl)
PknG (Ubl) (D188A)
PknG (Ubl) (E190A)
PknG (Ubl) (Q192A)
PknG (Ubl) (M194A)
PknG (Ubl) (M196A)
PknG (Ubl) (E198A)
PknG (Ubl) (R199A)
PknG (Ubl) (Q200A)
PknG (Ubl) (F201A)
PknG (Ubl) (L202A)
His-UbcH7
Mr (kDa)
IB: His
Input:
IB: GST
IB: His
17
11
35
D
GST-PknG
GST-PknG K181M
GST-PknG E190A
ATP
Mr (kDa)
p-Thr
input:
GST
135
100
E
GST
PknG (Ubl)
PknG (Ubl) E190A
Mr (kDa)
IB: His-UbcH7
Input:
IB: GST
IB: His-UbcH7
17
11
35
25

**Fig. S1.** Mtb PknG interacts with UbcH7 via Glu190 in the Ub-like (Ubl) domain. (*A*) Predicted region (red font) for UbcH7 interaction in the Ubl (green) domain of PknG (Left) and interfaces between Mtb PknG (green, ubiquitin-like domain; magenta, UbcH7 binding sites) and UbcH7 (orange) (Right). Mtb PknG PDB accession code: 2PZI; UbcH7 PDB accession code: 5UDH. (*B*) IP of GFP-tagged Mtb PknG or its truncated forms by Myc-UbcH7 in HEK293T cells. Red arrows indicate the bands of PknG-GFP, PknG (Ubl)-GFP, or PknG ΔUbl-GFP. (*C*) Pull-down of His-UbcH7 (2 μg each) by GST-tagged Ubl domain of PknG and its mutants (7 μg each). (*D*) Autophosphorylation level of PknG and its mutants *in vitro.* Autophosphorylation activity of recombinant GST-tagged PknG, PknG K181M, and PknG E190A was detected using anti-pThr antibody. Coomassie brilliant blue staining shows loading of the recombinant proteins. (*E*) Pull-down of His-tagged UbcH7 (3 μg each) by GST (6 μg), GST-tagged Mtb PknG (Ubl), or its E190A mutant (7 μg each).

### Figure S2

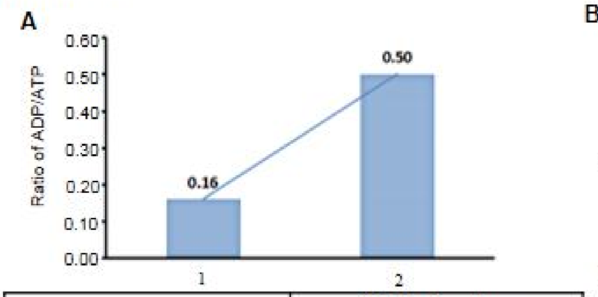


| | Samples | Relative abundance | | | |
|---|---|---|---|---|---|
| | | AMP | ADP | ATP | ADP/ATP |
| 1 | PknG K181M+ATP | ND | 1.71E+03 | 1.06E+04 | 0.16 |
| 2 | PknG K181M+ATP+UbcH7+Ub | ND | 4.74E+03 | 9.55E+03 | 0.5 |

ND: Not detected

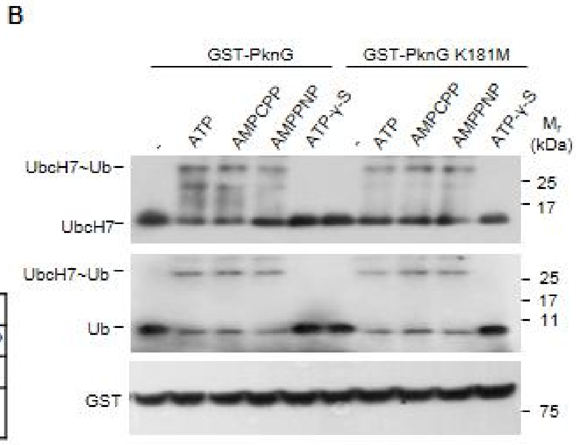


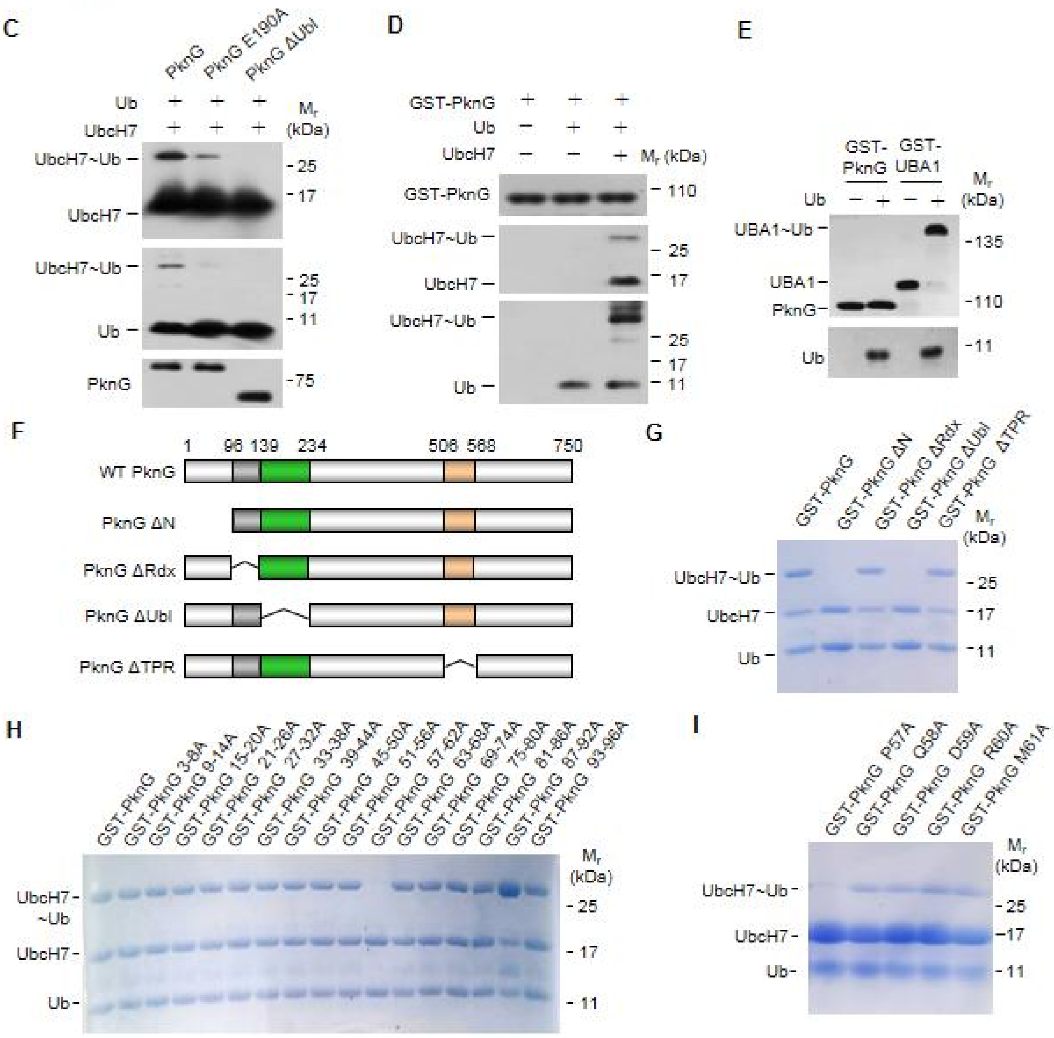

**Fig. S2.** Mtb PknG possesses unconventional Ub-activating enzyme activity. (*A*) The ADP/ATP ratios in the ubiquitination reaction mixtures. Reaction mixtures containing 2 μM GST-tagged PknG K181M, 5 μM ATP were incubated at 37 ℃ for 30 min in the presence (reaction 1) or absence (reaction 2) of 5 μM Ub and 5 μM UbcH7. LC-MS MultiQuant software was used to calculate the normalized abundance of each compound (including AMP, ADP, and ATP) by measuring the peak area intensity. (*B*) *In vitro* ubiquitin conjugation assay of UbcH7. UbcH7 was incubated with GST-tagged PknG or PknG K181M, and Ub at 37 ℃ for 30 min in the presence of ATP or ATP analogs including APCPP, AMPPNP, and ATP-γ-S. Reaction products were immunoblotted with antibodies against UbcH7 (top panel), Ub (middle panel) and the GST tag on PknG (bottom panel). (*C*) UbcH7 ubiquitin conjugation assay catalyzed by PknG or its mutants at 37 ℃ for 30 min. (*D*) UbcH7 ubiquitin conjugation assay *in vitro*. GST-tagged PknG was incubated with UbcH7 and Ub at 37 ℃ for 30 min in the presence of ATP. (*E*) *In vitro* ubiquitin conjugation assay of E1 (UBA1) and PknG. Reaction products were immunoblotted with antibodies against GST (top panel) and Ub (bottom panel). (*F*) Schematic diagram of Mtb PknG domains. PknG ΔN, N-terminal region (which was reported to be essential for Mtb intracellular survival)-deleted PknG; PknG ΔRdl, rubredoxin-like domain-deleted PknG; PknG ΔUbl: Ubiquitin-like domain-deleted PknG, PknG ΔTPR: TRP domain-deleted PknG. (*G*) UbcH7 ubiquitin conjugation assay *in vitro* by GST-tagged PknG or its truncations. (*H* and *I*) UbcH7 ubiquitin conjugation assay *in vitro* by GST-tagged PknG or its mutants (the residues covering the potential E1 activity sites were mutated into alanine).

**Figure S3**

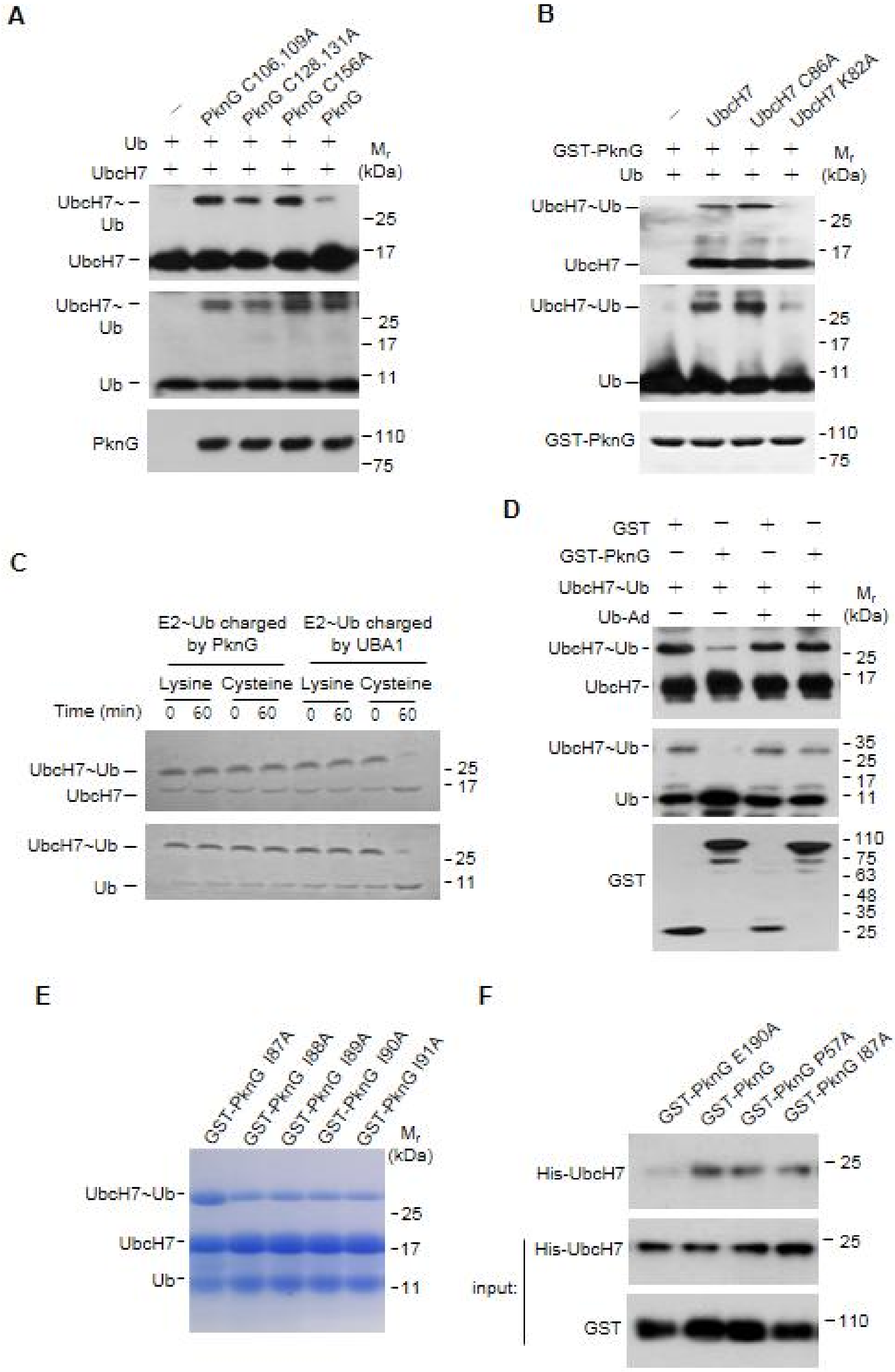
A
B
C
D
E
F
PknG C106,109A
PknG C128,131A
PknG C156A
PknG
Ub
UbcH7
UbcH7~Ub
PknG
Mr (kDa)
UbcH7 C86A
UbcH7 K82A
GST-PknG
E2~Ub charged by PknG
E2~Ub charged by UBA1
Lysine
Cysteine
Time (min)
GST
Ub-Ad
GST-PknG I87A
GST-PknG I88A
GST-PknG I89A
GST-PknG I90A
GST-PknG I91A
GST-PknG E190A
GST-PknG P57A
His-UbcH7
input:

**Fig. S3.** Mtb PknG possesses unconventional Ub-activating enzyme and isopeptidase activities. (*A*) UbcH7 ubiquitin conjugation assay catalyzed by PknG or its mutants at 37 ℃ for 30 min. Reactions were terminated with SDS sample buffer. (*B*) Ubiquitin conjugation assay of UbcH7 or its mutants as in (*A*). (*C*) *In vitro* ubiquitin discharge assay of UbcH7~Ub charged by PknG or E1 at 37 ℃ in buffers containing 50 mM free lysine or cysteine. (*D*) *In vitro* deubiquitinase assays of PknG with or without ubiquitin-aldehyde (Ub-Ad). Reactions were performed at 37 ℃ for 30 min and analyzed as in (*A*). (*E*) UbcH7 ubiquitin conjugation assay *in vitro* by GST-tagged PknG or its mutants (the residues covering the potential isopeptidase sites were mutated into alanines). (*F*) Pull-down of His-UbcH7 (2 μg each) by GST-tagged PknG, PknG P57A (an E1 activity-dead mutant), or PknG I87A (an isopeptidase activity-dead mutant) (10 μg each).

### Figure S4

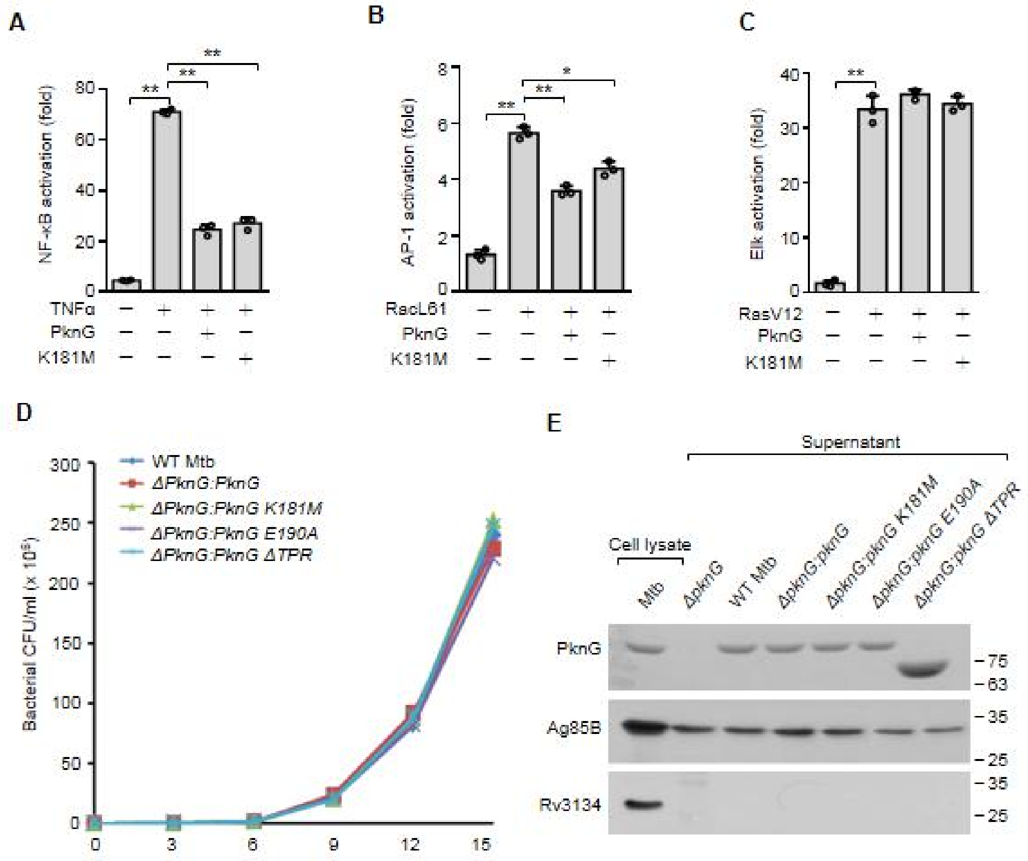

A
B
C
NF-κB activation (fold)
AP-1 activation (fold)
Elk activation (fold)
TNFα
RacL61
RasV12
PknG
K181M
D
Bacterial CFU/ml (x 10³)
WT Mtb
ΔPknG:PknG
ΔPknG:PknG K181M
ΔPknG:PknG E190A
ΔPknG:PknG ΔTPR
E
Supernatant
Cell lysate
Mtb
ΔpknG
WT Mtb
ΔpknG:pknG
ΔpknG:pknG K181M
ΔpknG:pknG E190A
ΔpknG:pknG ΔTPR
PknG
Ag85B
Rv3134

**Fig. S4.** Mtb PknG inhibits NF-κB signaling activation. (*A* to *C*) Luciferase assay of TNFα-induced NF-κB activation (*A*), RacL61-induced AP-1 activation (*B*), and RasV12-induced Elk activation (*C*) in the absence or presence of WT Mtb PknG or its K181M mutant form in HEK293T cells. Cells were transfected with the indicated plasmids for 24 h. (*D*) The *PknG* mutant strain exhibited no growth defect as compared to the WT Mtb strain. A total of $2 \times 10^6$ CFU of Mtb strains were inoculated into Middlebrook 7H9 broth and cultivated at 37 ℃, and then the culture from each time point were plated on 7H10 agar for bacterial CFU counting. (*E*) Immunoblotting analysis for protein levels of PknG from culture supernatants of the indicated Mtb strains. Cultures were grown in 50 ml Sauton medium for 2 days, and then the supernatants were collected for precipitation using 10% trichloroacetic acid. The secreted protein Ag85B was used as the loading control, and non-secreted protein Rv3134 was used to confirm that the supernatants were free of contamination due to cell lysis. Data are shown as mean ± SEM of three replicates. $*P < 0.05$ and $** P < 0.01$ (one-way ANOVA).

**Figure S5**

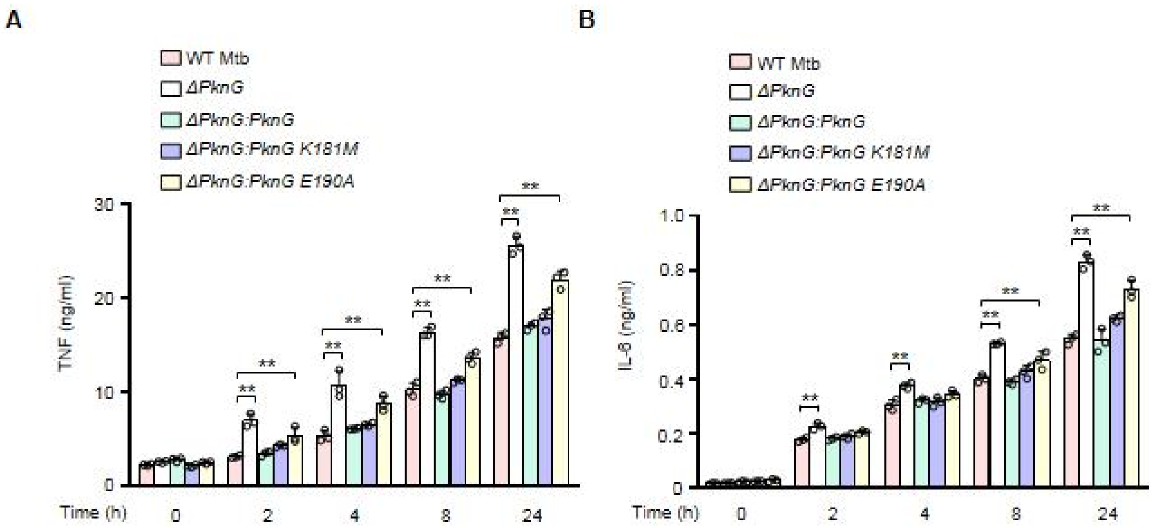
A
WT Mtb
ΔPknG
ΔPknG:PknG
ΔPknG:PknG K181M
ΔPknG:PknG E190A
TNF (ng/ml)
30
20
10
0
Time (h)
0
2
4
8
24
**
B
WT Mtb
ΔPknG
ΔPknG:PknG
ΔPknG:PknG K181M
ΔPknG:PknG E190A
IL-6 (ng/ml)
1.0
0.8
0.6
0.4
0.2
0
Time (h)
0
2
4
8
24
**

**Fig. S5.** Mtb PknG suppresses the production of TNF and IL-6 in macrophages. (*A* and *B*) ELISA of TNF (*A*) and IL-1β (*B*) in the medium of colony-stimulating factor-induced human primary monocyte-derived macrophages. Cells were infected with WT, *ΔpknG*, *ΔpknG:pknG*, *ΔpknG:pknG K181M* or *ΔpknG:pknG E190A* Mtb strain at an MOI of 1 for 0–24 h. Data are shown as mean ± SEM of three replicates. *$P < 0.05$ and ** $P < 0.01$ (one-way ANOVA).

**Figure S6**

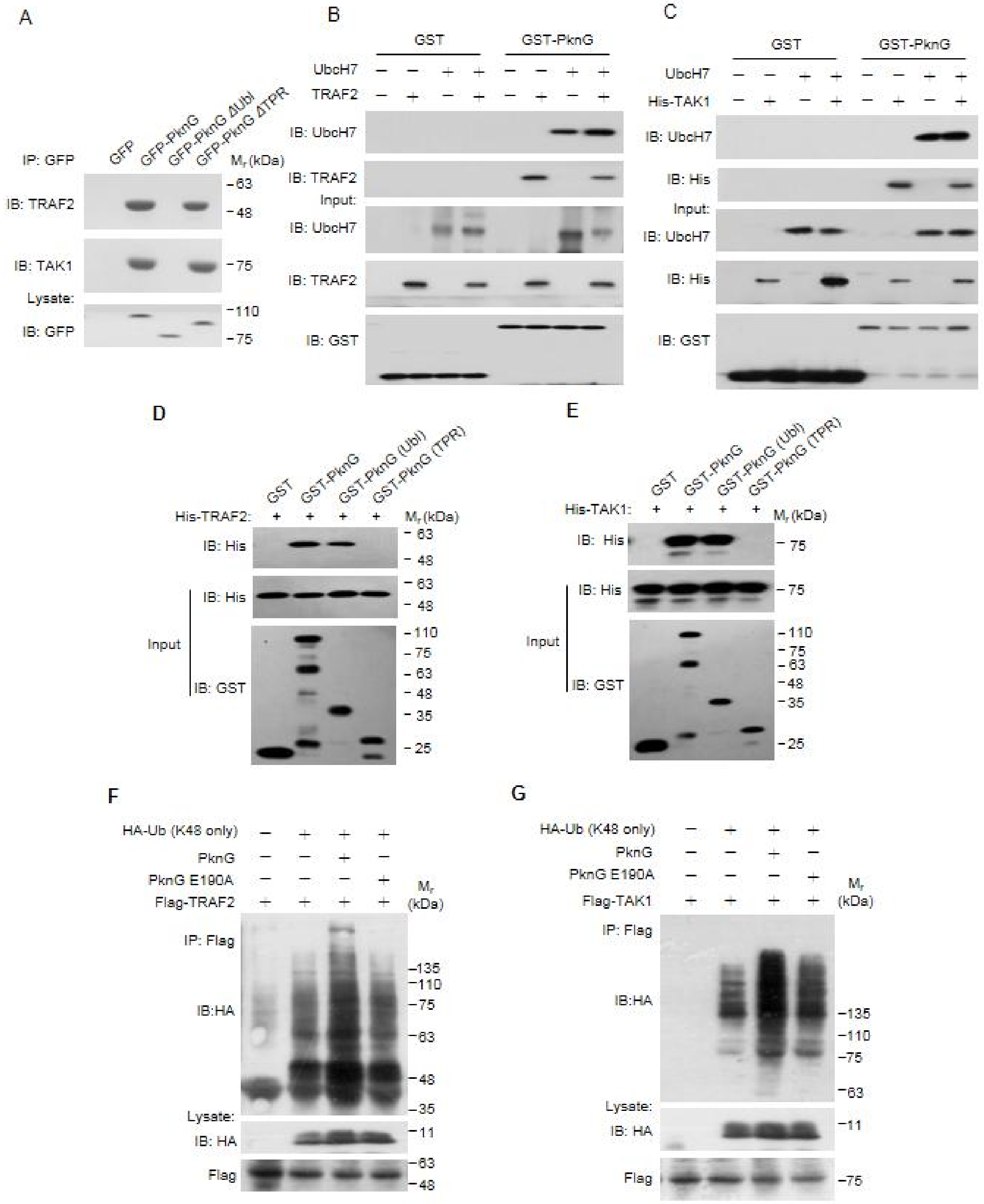


162

163

164

165

**Fig. S6.** Mtb PknG targets TRAF2 and TAK1 for degradation by binding to UbcH7. (*A*) Immunoblotting analysis of TRAF2 and TAK1 proteins immunoprecipitated using anti-GFP antibody from lysates of HEK293T cells transfected with GFP-tagged Mtb PknG or its truncated forms for 24 h. (*B* and *C*) Pull-down of TRAF2 (6 μg each) (*B*) or TAK1 (8 μg each) (*C*) by GST (3 μg each) or GST-tagged WT PknG (10 μg each) with or without purified UbcH7 (2 μg each). (*D* and *E*) Pull-down of His-tagged TRAF2 (6 μg each) (*D*) or TAK1 (8 μg each) (*E*) by GST-tagged WT PknG (10 μg each) or its truncated forms (3 μg each). (*F* and *G*) *In vivo* ubiquitination assay of TRAF2 (*F*) and TAK1 (*G*) in HEK293T cells. Cells were transfected with vectors encoding Flag-TRAF2, HA-Ub (K48 only), and GFP-tagged Mtb PknG or its E190A mutant form for 24 h.

**Table S1. Plasmids, bacterial strains and oligonucleotides used in this study.**

| **Name** | **Description** | **Reference** |
|---|---|---|
| **Plasmids** | | |
| pEGFP-N1 | CMV promoter, for mammalian expression, GFP tag, $Kan^R$ | Clontech |
| pEGFP-N1-PknG | For expression of GFP-PknG in mammalian cells | This study |
| pEGFP-N1-PknG K181M | For expression of GFP-PknG K181M in mammalian cells | This study |
| pEGFP-N1-PknG (Ubl) | For expression of GFP-PknG (Ubl) in mammalian cells | This study |
| pEGFP-N1-PknG (△Ubl) | For expression of GFP-PknG (△Ubl) in mammalian cells | This study |
| pEGFP-N1-PknG E190A | For expression of GFP-PknG E190A in mammalian cells | This study |
| pEGFP-N1-PknG ΔTPR | For expression of GFP-PknG ΔTPR in mammalian cells | This study |
| pEGFP-N1-PknG TPR | For expression of GFP-PknG TPR in mammalian cells | This study |
| p3xFlag-CMV14 | CMV promoter, for mammalian expression, 3xFlag tag, $Amp^R$ | Sigma |
| p3xFlag-CMV14-PknG | For expression of 3x Flag-PknG in mammalian cells | This study |
| p3xFlag-CMV14-PknG K181M | For expression of 3x Flag-PknG K181M in mammalian cells | This study |
| p3xFlag-CMV14-PknG E190A | For expression of 3x Flag-PknG E190A in mammalian cells | This study |
| p3xFlag-CMV14-PknG (Ubl) | For expression of 3x Flag-PknG (Ubl) in mammalian cells | This study |
| p3xFlag-CMV14-TRAF2 | For expression of 3x Flag-TRAF2 in mammalian cells | This study |
| p3xFlag-CMV14-TAK1 | For expression of 3x Flag-TAK1 in mammalian cells | This study |
| pcDNA6A | T7 promoter, for mammalian expression, Myc tag, $Amp^R$ | Invitrogen |
| pcDNA6A-UbcH7 | For expression of Myc-UbcH7 in mammalian cells | This study |
| pGEX-6P-1-UbcH7 | For expression of GST-UbcH7 in mammalian cells | This study |
| pGEX-6P-1-UbcH7 C86A | For expression of GST-UbcH7 C86A in mammalian cells | This study |
| pGEX-6P-1-UbcH7 K72A | For expression of GST-UbcH7 K72A in mammalian cells | This study |
| pGEX-6P-1-UbcH5a | For expression of GST-UbcH5a in mammalian cells | This study |
| pGEX-6P-1-UbcH5b | For expression of GST-UbcH5b in mammalian cells | This study |
| pGEX-6P-1-UbcH5c | For expression of GST-UbcH5c in mammalian cells | This study |
| pGEX-6P-1-UbcH8 | For expression of GST-UbcH8 in mammalian cells | This study |
| pET30a | T7 promoter, for bacterial expression, 6xHis tag, $Kan^R$ | Novagen |
| pET30a-PknG | For expression of recombinant protein $His_6$-PknG | This study |
| pET30a-Ub | For expression of recombinant protein $His_6$-Ub | This study |
| pET30a-UbcH7 | For expression of recombinant protein $His_6$-UbcH7 | This study |
| pET30a-E1 | For expression of recombinant protein $His_6$-E1 | This study |
| pGEX-6P-1 | *tac* promoter, for bacterial expression, GST tag, $Amp^R$ | GE |
| pGEX-6P-1-PknG | For expression of recombinant protein GST-PknG | This study |
| pGEX-6P-1-PknG C106, 109A | For expression of recombinant protein GST-PknG C106,109A | This study |
| pGEX-6P-1-PknG C128, 131A | For expression of recombinant protein GST-PknG C128,131A | This study |
| pGEX-6P-1-PknG C156A | For expression of recombinant protein GST-PknG C156A | This study |
| pGEX-6P-1-PknG E190A | For expression of recombinant protein GST-PknG E190A | This study |
| pGEX-6P-1-PknG K181M | For expression of recombinant protein GST-PknG K181M | This study |
| pGEX-6P-1-PknG P57A | For expression of recombinant protein GST-PknG P57A | This study |
| pGEX-6P-1-PknG I87A | For expression of recombinant protein GST-PknG I87A | This study |
| pGEX-6P-1-PknG (TPR) | For expression of recombinant protein GST-PknG (TPR) | This study |

| | | |
|---|---|---|
| pGEX-6P-1-PknG (Ubl) E190A | For expression of recombinant protein GST-PknG (Ubl) E190A | This study |
| pGEX-6P-1-PknG ΔN | For expression of recombinant protein GST-PknG ΔN | This study |
| pGEX-6P-1-PknG ΔRdx | For expression of recombinant protein GST-PknG ΔRdx | This study |
| pGEX-6P-1-PknG ΔTPR | For expression of recombinant protein GST-PknG ΔTPR | This study |
| pGEX-6P-1-PknG ΔUbl | For expression of recombinant protein GST-PknG ΔUbl | This study |
| pGEX-6P-1-UBA1 | For expression of recombinant protein GST-UBA1 | This study |
| pcDNA3-HA-Ub | For expression of HA-Ub in mammalian cells | F. Shao |
| pcDNA3-HA-Ub (K48 only) | For expression of HA-Ub (K48 only) in mammalian cells | F. Shao |
| pcDNA3-HA-Ub (K63 only) | For expression of HA-Ub (K63 only) in mammalian cells | F. Shao |
| pNF-κB-luc | Used in dual-luciferase assay for NF-κB pathway | F. Shao |
| pRL-TK | Used in dual-luciferase assay for NF-κB and MAPK pathways | F. Shao |
| pGal4-luc | Used in dual-luciferase assay for MAPK pathway | F. Shao |
| pGal4-Elk | Used in dual-luciferase assay for Erk pathway | F. Shao |
| pFA-cJun | Used in dual-luciferase assay for MAPK pathway | F. Shao |
| pcDNA3-RacL61 | For expression of constitutively activated Rac in mammalian cells | F. Shao |
| pcDNA3-HA-RasV12 | For expression of RasV12 in mammalian cells | F. Shao |
| pCS2-3xFlag-TRAF2 | For expression of Flag-TRAF2 in mammalian cells | F. Shao |
| pCS2-3xFlag-TAB1 | For expression of Flag-TAB1 in mammalian cells | F. Shao |
| pCS2-3xFlag-TAB2 | For expression of Flag-TAB2 in mammalian cells | F. Shao |
| pCS2-3xFlag-TAB3 | For expression of Flag-TAB3 in mammalian cells | F. Shao |
| pCS2-3xFlag-TAK1 | For expression of Flag-TAK1 in mammalian cells | F. Shao |
| pGADT7 | Used for expressing proteins fused to GAL4 activation domain (AD) in the yeast, $Amp^R$ | This study |
| pGBKT7 | Used for expressing proteins fused to GAL4 DNA-binding domain (BD) as a bait expression vector in the yeast, $Kan^R$ | This study |
| pGADT7-UbcH7 | For expression of AD-UbcH7 in the yeast | This study |
| pGBKT7-PknG | For expression of BD-PknG in the yeast | This study |
| pGADT7-TAK1 | For expression of AD-TAK1 in the yeast | F. Shao |
| pGBKT7-TAB2 | For expression of BD-TAB2 in the yeast | F. Shao |
| pJV53 | Used to create *pknG* deletion mutant in mycobacteria; Expresses the mycobacteriophage Che9c gp60 and gp61 under acetamide induction, oriC, oriM, $Kan^R$ | Addgene |
| pYUB854 | Used for cloning allelic exchange substrate; Contains λ phage *cos* site, $Hyg^r$ | W. R. Jacobs |
| pYUB854-PknG KO | pYUB854 with *pknG*-disrupted sequences for generating Δ*pknG* | This study |
| pMV306 | Integrative vector used to complement the strain Mtb Δ*pknG*; Cloning vector replicating in *E. coli* with the kanamycin resistance gene *aph* from transposon Tn903 and the gene for the integrase and the *attP* site of phage L5 for integration into the mycobacterial genome | W. R. Jacobs |
| pMV306-PknG | pMV306 carrying the *pknG* gene and promoter for Mtb Δ*pknG* | This study |

| | | |
|---|---|---|
| | complementation | |
| pMV306-PknG K181M | pMV306 carrying the *pknG K181M* gene and promoter for creating mutant strain Mtb (pknG K181M) | This study |
| pMV306-PknG E190A | pMV306 carrying the *pknG E190A* gene and promoter for creating mutant strain Mtb pknG E190A | This study |
| pMV306-PknG ΔTPR | pMV306 carrying the *pknG ΔTPR* gene and promoter for creating mutant strain Mtb pknG ΔTPR | This study |
| **Bacterial strains** | | |
| *E. coli* DH5α | $F^-$ φ80*lac*ZΔM15 Δ (*lac*ZYA-*arg*F) U169 *rec*A1 *end*A1 *hsd*R17 (rk-, mk+) *pho*A *sup*E44 $\lambda^-$*thi*$^-$1 *gyr*A96 *rel*A1 | Invitrogen |
| *E. coli* BL21 (DE3) | $F^-$ *ompT hsdS*$_B$ ($r_B^-$ $m_B^-$) *gal dcm* (DE3) | Novagen |
| *M. tuberculosis* H37Rv | ATCC 27294 | ATCC |
| *Mtb ΔpknG* | Mtb strain with deletion of *pknG* | This study |
| *Mtb ΔpknG:pknG* | Created by introducing the integrative vector pMV306 carrying the *pknG* gene and promoter into the Mtb Δ*pknG* strain | This study |
| *Mtb ΔpknG:pknG K181M* | Created by introducing the integrative vector pMV306 carrying the *pknG K181M* gene and promoter into the Mtb Δ*pknG* strain | This study |
| *Mtb ΔpknG:pknG E190A* | Created by introducing the integrative vector pMV306 carrying the *pknG E190A* gene and promoter into the Mtb Δ*pknG* strain | This study |
| *Mtb ΔpknG:pknG ΔTPR* | Created by introducing the integrative vector pMV306 carrying the *pknG ΔTPR* gene and promoter into the Mtb Δ*pknG* strain | This study |
| **Oligonucleotides (5'-3')** | | |
| *Gapdh*-QRT-F | GGAGCGAGATCCCTCCAAAAT | This study |
| *Gapdh*-QRT-R | GGCTGTTGTCATACTTCTCATGG | This study |
| *Tnf*-QRT-F | CCTCTCTCTAATCAGCCCTCTG | This study |
| *Tnf*-QRT-R | GAGGACCTGGGAGTAGATGAG | This study |
| *Il6*-QRT-F | ACTCACCTCTTCAGAACGAATTG | This study |
| *Il6*-QRT-R | CCATCTTTGGAAGGTTCAGGTTG | This study |
